# The nucleocytoplasmic translocation of HINT1 regulates the maturation of cell density

**DOI:** 10.1101/2025.01.13.632869

**Authors:** Xiaofang Zhang, Fumihiko Nakamura

## Abstract

Cell contact inhibition progresses through three stages: (i) increasing cell density leads to reduced movement while mitosis continues; (ii) a rapid shift to epithelial morphology; and (iii) ongoing division with decreased cell size. This transition involves reorganizing the actin cytoskeleton from stress fibers to a cortical network, stabilizing cell shape, and strengthening cell-cell connections. However, the signaling pathways regulating the final stage remain unclear. We identified histidine triad nucleotide-binding protein 1 (HINT1), also known as protein kinase C inhibitor 1 (PKCI-1), as essential for monolayer maturation. At low density, HINT1 is located in the nucleus, binding to open chromatin. As density increases, exportin-1 relocates HINT1 to the cytoplasm, where it inhibits PKC and remodels the actin cytoskeleton. While monolayer formation can occur without HINT1, its presence is necessary for fully confining cells and achieving a mature monolayer. Additionally, MARCKS phosphorylation decreases in high-density cells, and loss of HINT1 leads to increased cell area, similar to hypertrophic conditions in patients with low HINT1 levels.

**Summary blurb:** Histidine triad nucleotide-binding protein 1 shuttles between the nucleus and cytoplasm based on cell density, regulating gene expression in the nucleus and inhibiting protein kinase C in the cytoplasm to remodel actin.

## Introduction

Early unicellular organisms evolved to proliferate indefinitely under favorable environmental conditions, such as sufficient nutrient availability. In contrast, multicellular organisms developed regulatory mechanisms that suppress unchecked cell division, thus enabling the formation of distinct body architectures. Cancer arises when these regulatory mechanisms fail, resulting in the uncontrolled expression of intrinsic proliferative capacities.

Such inhibitory mechanisms are known as contact inhibition of proliferation (CIP) [1, 2]. CIP proceeds through three distinct stages: (i) a phase of increasing cell density during which cell motility declines (contact inhibition of locomotion, CIL) but mitosis persists; (ii) a rapid transition to an epithelial morphology; and (iii) continued cell division that progressively reduces cell size (cell density maturation) and slows the mitotic rate. Ultimately, mitosis ceases once the cell area drops below a critical threshold [3]. These observations suggest that CIP arises from mechanical interactions and constraints rather than simple cell-cell contact. However, the molecular mechanisms that regulate these processes, particularly the final stage, remain largely unexplored.

During these transitions, the actin cytoskeleton undergoes remodeling. When tissue culture cells are grown at low densities, these cells migrate and frequently exhibit polarized stress fibers (SFs). As cell density increases during high-density culture, SFs tend to disappear, and filamentous actin (F-actin) is redistributed to the cell cortex, resembling its *in situ* distribution [4–10]. These arrangements suppress cell spreading and movement and support maintenance of cell shape and reinforcement with neighboring cells.

During CIL and CIP, various molecules, such as cadherins, focal adhesion kinase, and small GTPases remodel the cytoskeleton to promote monolayer formation [1, 2, 11–14]. Our findings reveal that histidine triad nucleotide-binding protein 1 (HINT1), also known as protein kinase C inhibitor 1 (PKCI-1) [15, 16], plays a role in further maturing cells within the monolayer, leading to smaller cell shapes. HINT1, a tumor suppressor gene, is involved in various functions such as cell signaling and gene expression [17]. For example, HINT1 acts as a transcriptional co-regulator, influencing gene expression by interacting with various transcription factors such as MITF (microphthalmia-associated transcription factor) and β-catenin to regulate cell proliferation, apoptosis, and differentiation [18]. Here, we show that at low cell density, HINT1 predominantly resides in the nucleus and binds to chromatin. However, as cell density increases, independent of cell-cell attachment, HINT1 translocates to the cytoplasm, where it disassembles SFs by inhibiting PKC. Supporting this model, forced expression of HINT1 in the cytoplasm, achieved by mutating its nuclear localization signal (NLS), reduces SFs and induces rounded cell morphologies instead of extended ones. Similarly, pharmacological inhibition of exportin 1 traps HINT1 in the cytoplasm, also resulting in rounded cell shapes. Additionally, loss of HINT1 increases cell cross-sectional area at high density, consistent with hypertrophy observed in cardiomyocytes from HINT1 knock-out mice and in hypertrophic patients expressing low levels of HINT1, though the molecular mechanism was previously unknown. These findings establish HINT1 as a crucial molecule that shuttles between the nucleus and cytoplasm in a cell-density-dependent manner. Regulated by active transport at high cell densities, HINT1 promotes mature cell layering by remodeling actin filaments through the inhibition of protein kinase C. Consequently, our findings elucidate previous reports that CIP progresses through three distinct phases, driven by HINT1-mediated signaling pathways operating in both the nucleus and cytoplasm.

## Results

### HINT1 is a cell density-dependent nucleocytoplasmic shuttling molecule

We recently developed a DSP-MNase-proteogenomics method to identify proteins that bind to open chromatin [19]. HINT1 was identified as a potential chromatin-binding protein in low-density human skeletal muscle (hsSKM) cells, with reduced binding observed in high-density cells (Table S1 in ref. [19]). In this study, we confirmed whether HINT1 can shuttle between the nucleus and cytoplasm in a cell density-dependent manner. First, we expressed exogenous HINT1 with a hemagglutinin (HA) tag attached to the C-terminus in HEK293A cells at both low and high densities (**Fig. 1A and 1B**). Consistent with the proteomics data, HINT1-HA was primarily localized in the nucleus at low density, while it was found in the cytoplasm at high density. To visualize the localization of endogenous HINT1in various tissue culture cells, we used a commercially available specific antibody (**Fig. 1C**). Cell density did not affect the expression level of HINT1 or induce fragmentation. Consistent with the results for exogenously expressed HINT1-HA, endogenous HINT1 was also primarily localized in the nucleus of HEK293A, hsSKM, and mouse embryonic fibroblast (MEF) cells at low density, with slight variations depending on the cell type (**Fig. 1D and 1E**). At high density, however, HINT1 predominantly relocated to the cytoplasm.

**Fig. 1.**
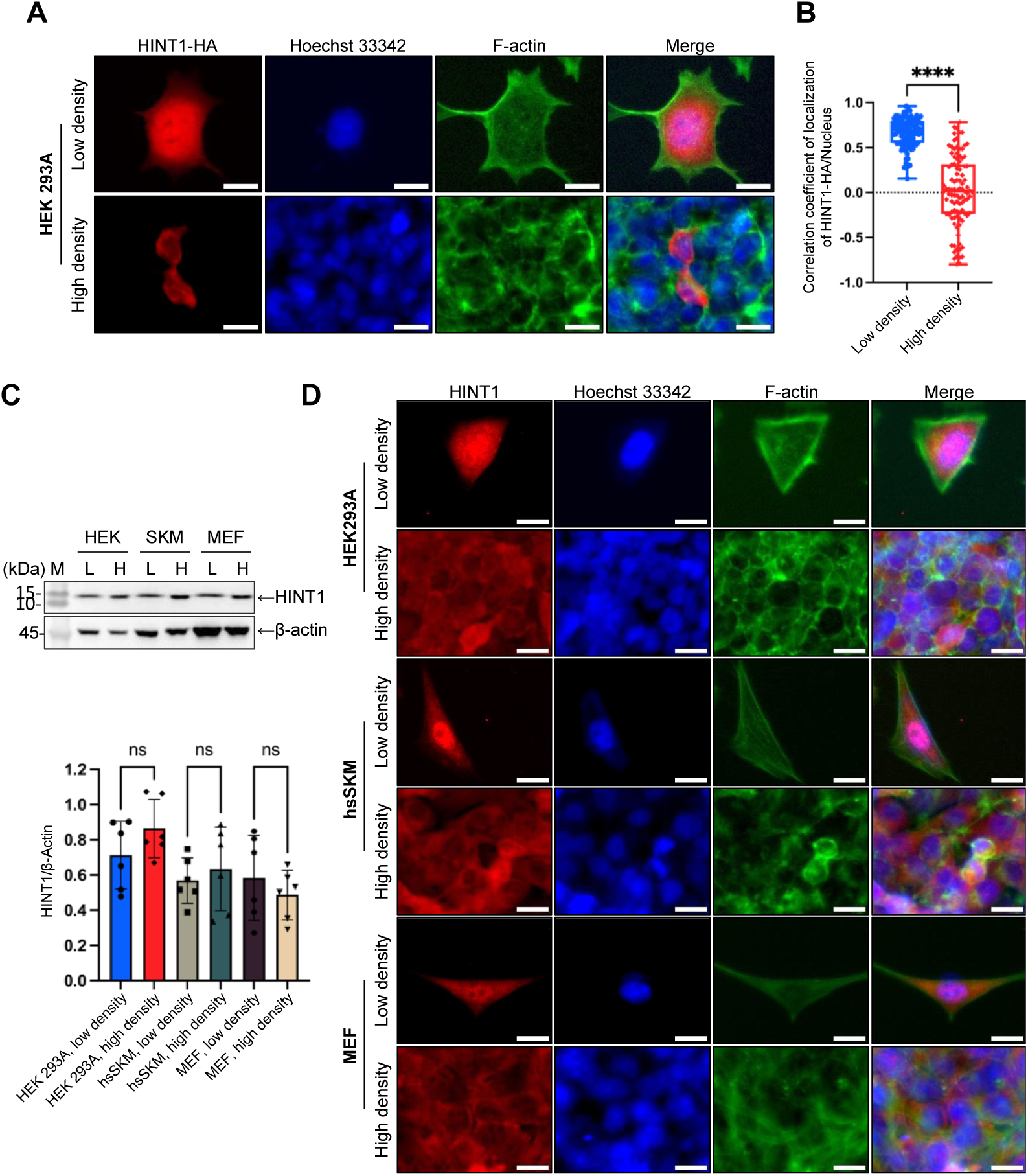
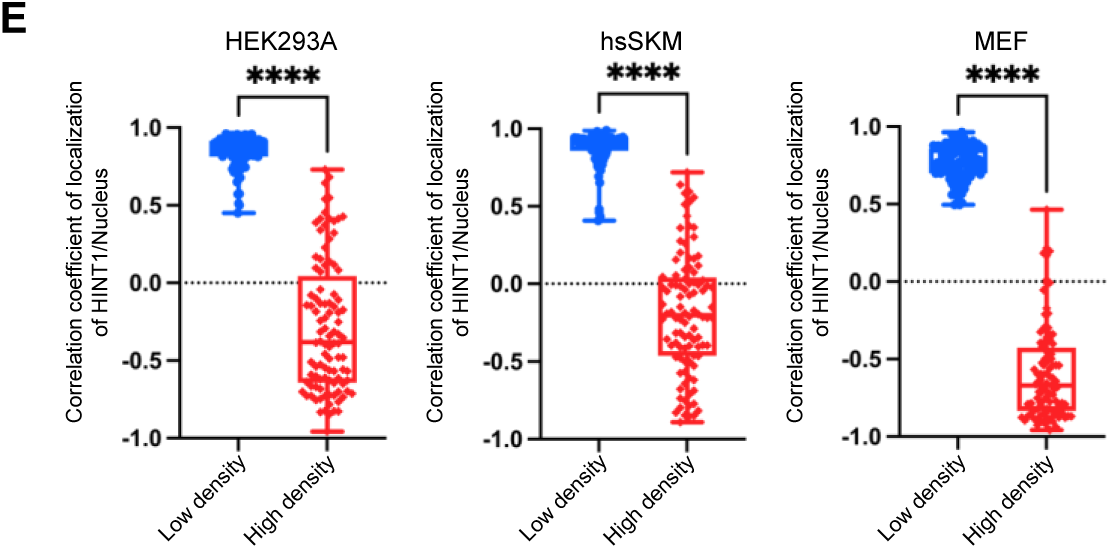
HINT1 translocates between the nucleus and cytoplasm in a cell density-dependent manner (A) Exogenously expressed HINT1-HA in HEK293A cells at low density mainly localizes in the nucleus, whereas it translocates to the cytoplasm at high density. HINT1-HA was stained with rabbit anti-HA antibody followed by anti-rabbit IgG- Alexa Fluor Plus 594 antibody (red). Nuclear DNA was stained with Hoechst 33342 (blue). Actin filaments were stained with phalloidin Alexa Fluor 488 (green). Scale: 20 μm. (B) Correlation coefficient of HINT1-HA staining and Hoechst nuclear staining were calculated and plotted (n=100). (C) Expression of HINT1 in various tissue culture cells detected by western blotting using specific antibody against HINT1. (D) Localization of endogenous HINT1 expressed in various tissue culture cells at low and high densities. HINT1 was stained with rabbit anti-HINT1 antibody followed by anti-rabbit IgG- Alexa Fluor Plus 594 antibody. Nuclear DNA was stained with Hoechst 33342. Actin filaments were stained with phalloidin Alexa Fluor 488. Scale: 20 μm. (E) Correlation coefficient of HINT1 staining and Hoechst 33342 staining were calculated and plotted (n=100).

Cell-cell contact can induce contact inhibition of proliferation (CIP) through various transmembrane proteins [1, 2]. However, simple cell confluency is not sufficient to trigger YAP translocation to the cytoplasm; much higher cell density is required for this process to occur [20]. While HINT1 remains in the nucleus when cells are in partial contact (e.g., 70% confluency in **Fig. S1**), near full confluency (e.g., 100%) is sufficient to initiate its translocation to the cytoplasm. As cell density increases, a greater amount of HINT1 relocates to the cytoplasm (**Fig. S1**).

### HINT1 is not mechanosensitive nucleocytoplasmic shuttling molecule

Many nucleocytoplasmic molecules, such as YAP1 and CBFB, display both density-dependent localization and mechanosensitivity, which is influenced by factors such as actin-myosin contraction and substrate stiffness [2, 21, 22]. To determine whether HINT1 shares this sensitivity to internal mechanical stress, we treated cells with latrunculin B (an actin polymerization inhibitor) and blebbistatin (a myosin II inhibitor). Unexpectedly, HINT1 translocation was unaffected by these treatments in various tissue culture cells (**Fig. 2A, 2B, and S2**), while YAP, as expected, translocated to the cytoplasm (**Fig. S3**), consistent with prior findings [21]. We also assessed HINT1 translocation in cells cultured on a soft substrate, but even on a 0.2 kPa substrate, HINT1 did not move to the cytoplasm (**Fig. 2C**).

**Fig. 2.**
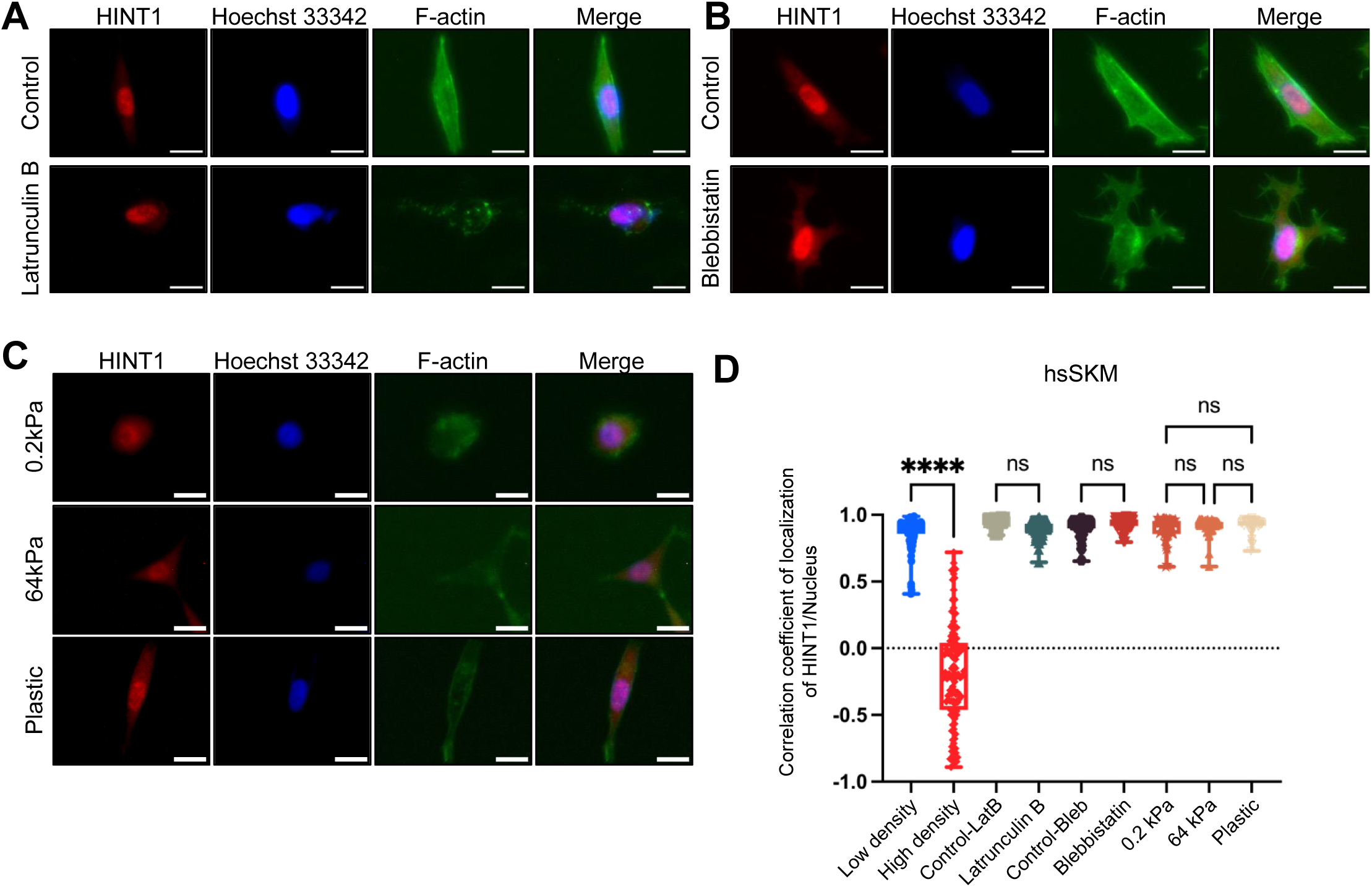
HINT1 is not mechanosensitive nucleocytoplasmic shuttling molecule (**A**∼**C**) cells were stained as shown in Fig. 1D. Scale: 20 μm. (A) hsSKM cells were treated with or without 5 μM of Latrunculin B for 60 min. (B) The cells were treated with or without 50 μM of blebbistatin for 180 min. (C) The cells were cultured on soft (0.2 kPa) and stiff substrate (64 kPa and plastic). (D) Correlation coefficient of HINT1 staining and Hoechst 33342 staining were calculated and plotted (In the 0.2 kPa, 64 kPa, and plastic groups, n=50; while in the other group, n=100). Related to Fig. S2.

### Loss of HINT1 delay cell density-dependent cell death but not cell migration

To explore the function of HINT1, we generated HINT1 knock-out (KO) HEK293A cells using the CRISPR- Cas9 system. Western blotting and immunofluorescence microscopy verified that three cloned KO cells lack HINT1 expression (**Fig. 3A and 3B**). Unexpectedly, no significant morphological changes were observed following the knockout (**Fig. 3B**). Next, we examined whether the HINT1 knockout affects cell migration (**Fig. 3C-3E**). Once again, no significant changes were observed. When each cell was cultured with daily fresh medium, no significant difference in proliferation was observed for the first 10 days. However, WT cells died after 6 days of reaching a peak. Interestingly, KO cells survived longer than WT cells (**Fig. 3F**).

**Fig. 3.**
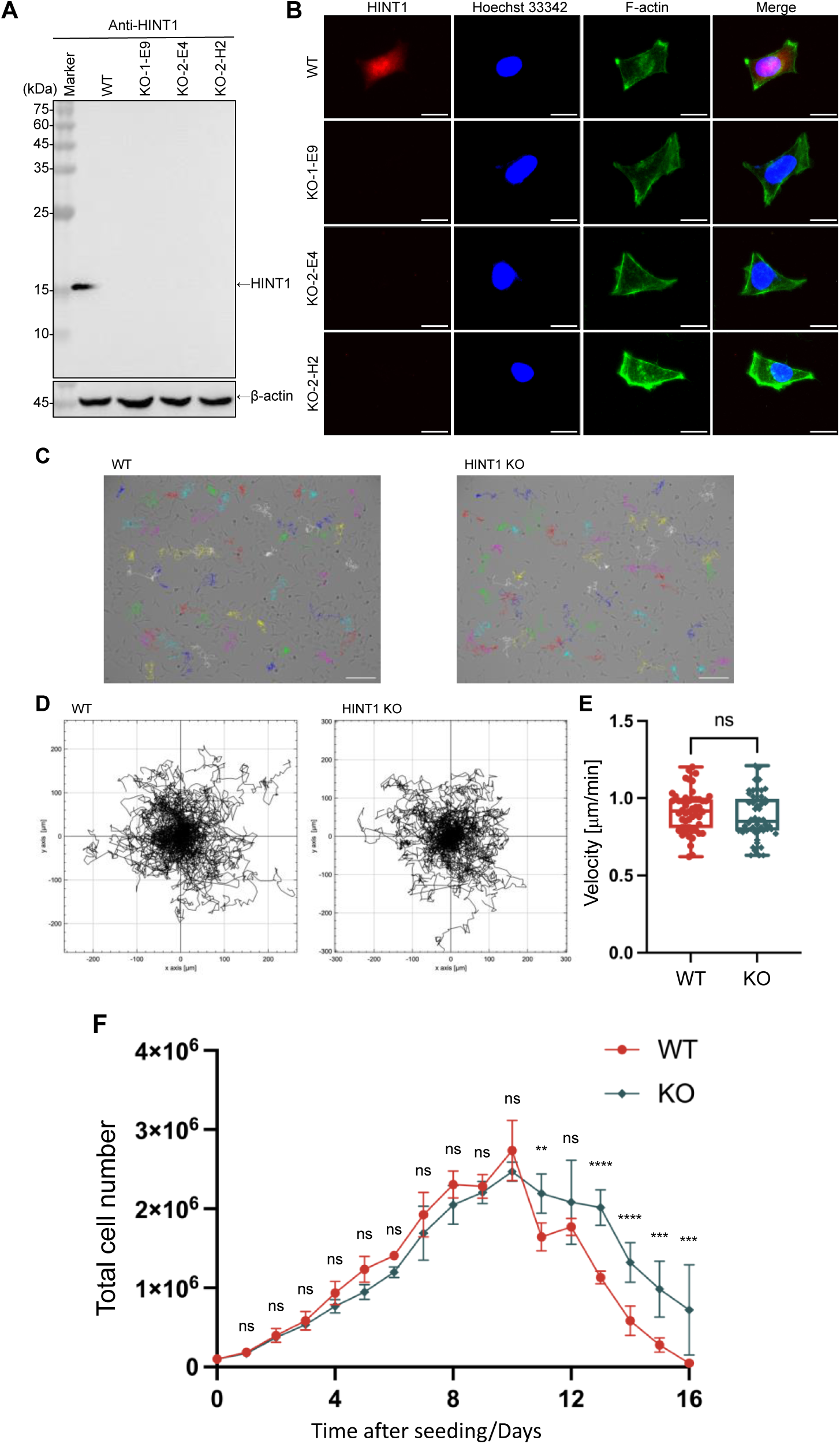
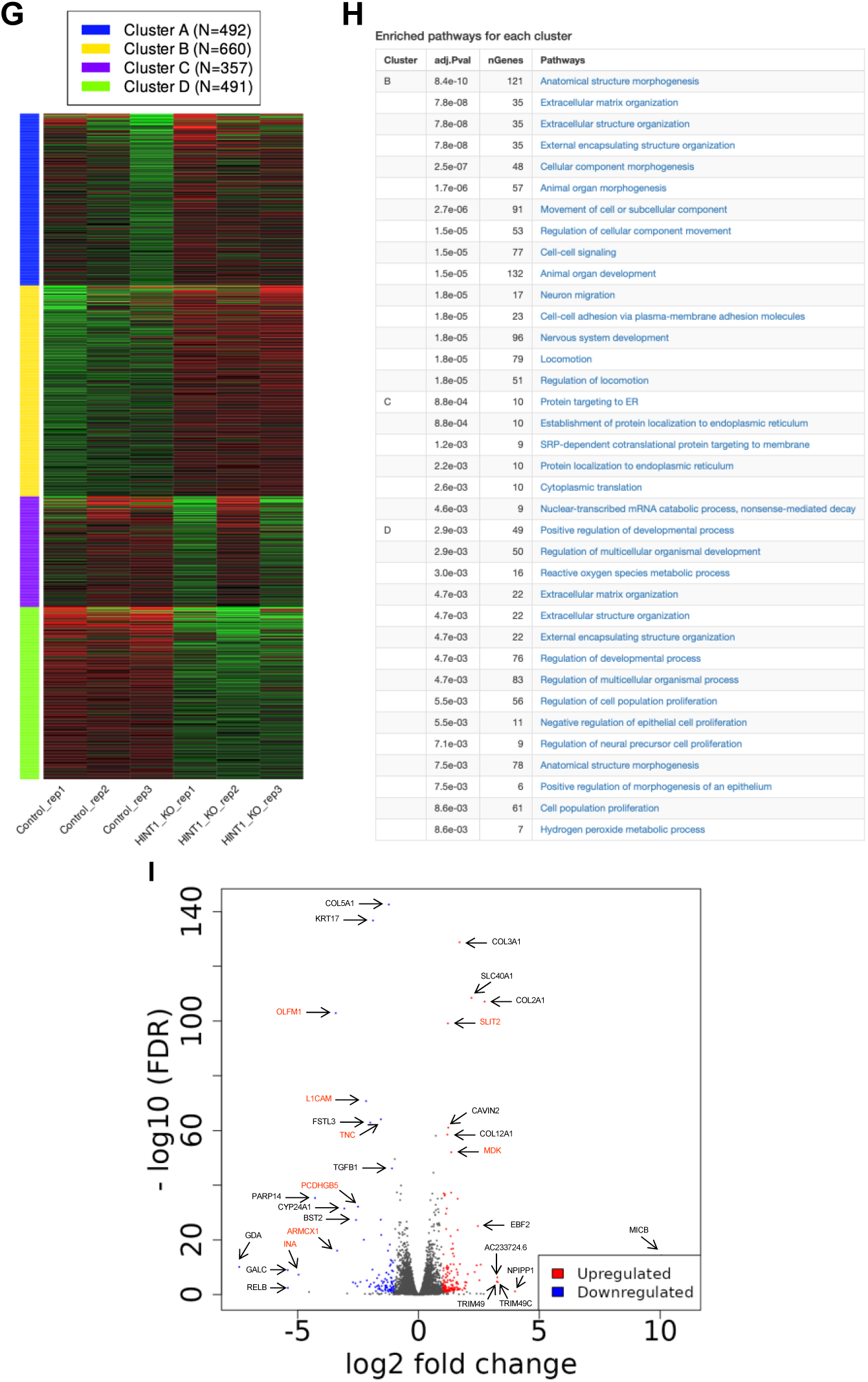
Loss of HINT1 delay cell density-dependent cell death (**A** and **B**) Western blotting and immunofluorescent microscopy confirm loss of HINT1 in three independent HINT1-KO cell lines. The cells were stained as shown in Fig. 1D. Scale: 20 μm. (**C**-**E**) Images of cells were captured at 1 frame/10 min for 20 h by time-lapse microscopy and tracked the migratory paths of cells. Scale: 200 μm. (D) Plots show an example tracking 50 cells from one of three independent experiments. (E) Mean migration speed (μm/min) of cells. Values represent the migration speed of each cell as a dot. Statistically not significant (ns) by the two-tailed unpaired Student’s test. (F) Equal number of WT and HINT1-KO HEK293A cells were cultured in 24-well tissue culture plate and medium were replaced every day. At the indicated time after seeding, cell number were counted and plotted. Average of three independent experiments. Statistical analysis was performed by two-way ANOVA. ns, not significant; * p≦0.05; ** p≦0.01, *** p≦0.001, **** p≦0.0001. (**G** and **H**) DEGs between WT and HINT1-KO cells. K-means clustering (The red and blue represent significantly up-and down-regulated genes between WT and HINT1-KO cells, respectively) and enrichment analysis. (**I**) The Volcano plot of false discovery rate (-log10 FDR) against log 2 (fold change). The red and blue dots represent significantly up-and down-regulated genes between WT and HINT1-KO cells, respectively. The genes indicated in red are involved in neural development.

Finally, we analyzed differentially expressed genes (DEGs) between WT and HINT1-KO cells using RNA-seq analysis. The results showed that HINT1 KO led to both the upregulation and downregulation of genes associated with morphogenesis and development (**Fig. 3G, 3H, and S4**). Although the Volcano plot highlighted significant up- and down-regulated genes between WT and HINT1-KO cells, with the upregulated genes being primarily involved in neural development (**Fig. 3I**), potentially linked to *HINT1* gene polymorphisms in human behaviors and neuropathy [23, 24], the RNA-seq analysis did not provide clear insights into specific biological functions.

### HINT1 regulates cell cross-sectional area at high density

Although no significant differences in cell morphology, proliferation, or migration were observed between WT and HINT1 KO cells at low density, KO cells survived longer than WT cells at high density, suggesting that HINT1 functions under high-density conditions. First, we stained actin filaments in WT and KO cells at various densities (**Fig. S5**). While no significant differences in actin distribution were observed between WT and KO cells, we noticed that the cell area of KO cells appeared larger than that of WT cells. To confirm this observation, cell boundaries were demarcated using a fluorogenic membrane dye to quantitatively assess cell cross-sectional area (**Fig. 4**). Consistent with the actin staining images, cross-sectional area of KO cells at high density is larger than that of WT cells, whereas at low density such difference was not observed.

**Fig. 4.**
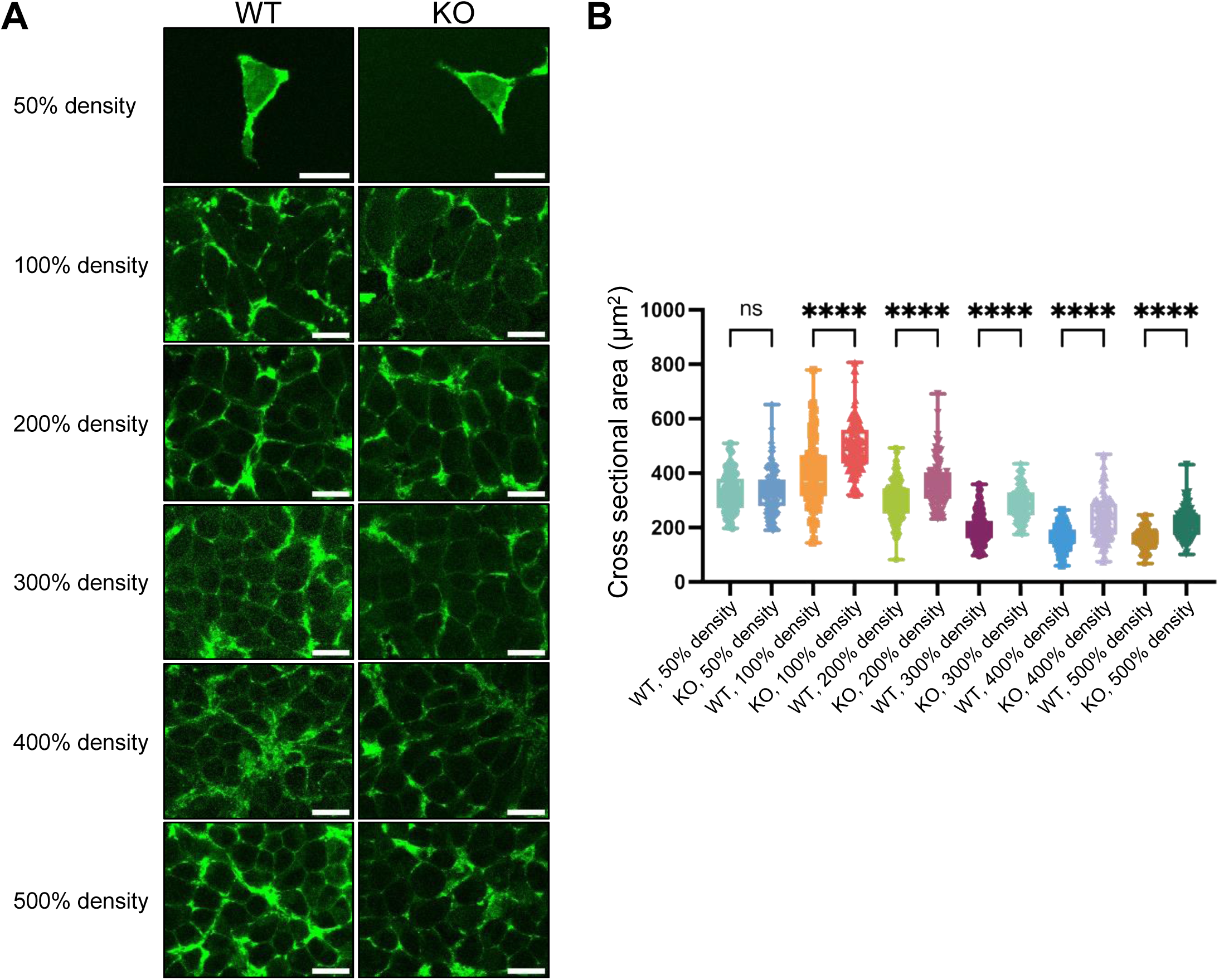
HINT1 KO enlarge cell cross-sectional area at high density (A) WT and HINT1-KO HEK293A cells were stained with fluorogenic membrane dye to demarcate the cell boundaries at various cell density. Scale: 20 μm. (B) Values represent the cross-sectional area of each cell as a dot (n=100). The percentages are values used for convenience when seeding cells and do not represent the actual cell density. However, for 100%, the cell count was measured and cultured in such a way that the cells would reach confluency by the time of fixation.

### Forced cytoplasmic expression of HINT1 disrupts actin SF and promotes a rounded, confined cell morphology

To investigate molecular mechanism of how translocation of HINT1 to the cytoplasm regulate cell morphology at high density, we engineered HINT1 that are expected to be constitutively expressed in the nucleus or cytoplasm. Although attachment of nuclear localization signal (NLS, MPKKKRKV-) to the N-terminal of HINT1 promote translocation of HINT1 to the nucleus in low density cells, the engineered HINT1 still stay in the cytoplasm at high density (**Fig. S6 and S7**). A mutation in the NLS sequence (MPK<u>T</u>KRKV-), which is expected to disrupt nuclear localization [25], only slightly reduced the NLS’s effect on nuclear translocation at low density, without significantly altering the overall intracellular distribution of HINT1. However, point mutation of the predicted NLS (**Fig. S6D**) in the HINT1 molecule disrupted the nuclear localization as expected (**Fig. 5A and 5B**). Furthermore, in MEF cells, the exogenously expressed HINT1-HA with the R24A/R25A mutation impaired actin SF and significantly altered cell morphology, causing the cells to become smaller, rounder, and more compact. (**Fig. 5C and 5D**).

**Fig. 5.**
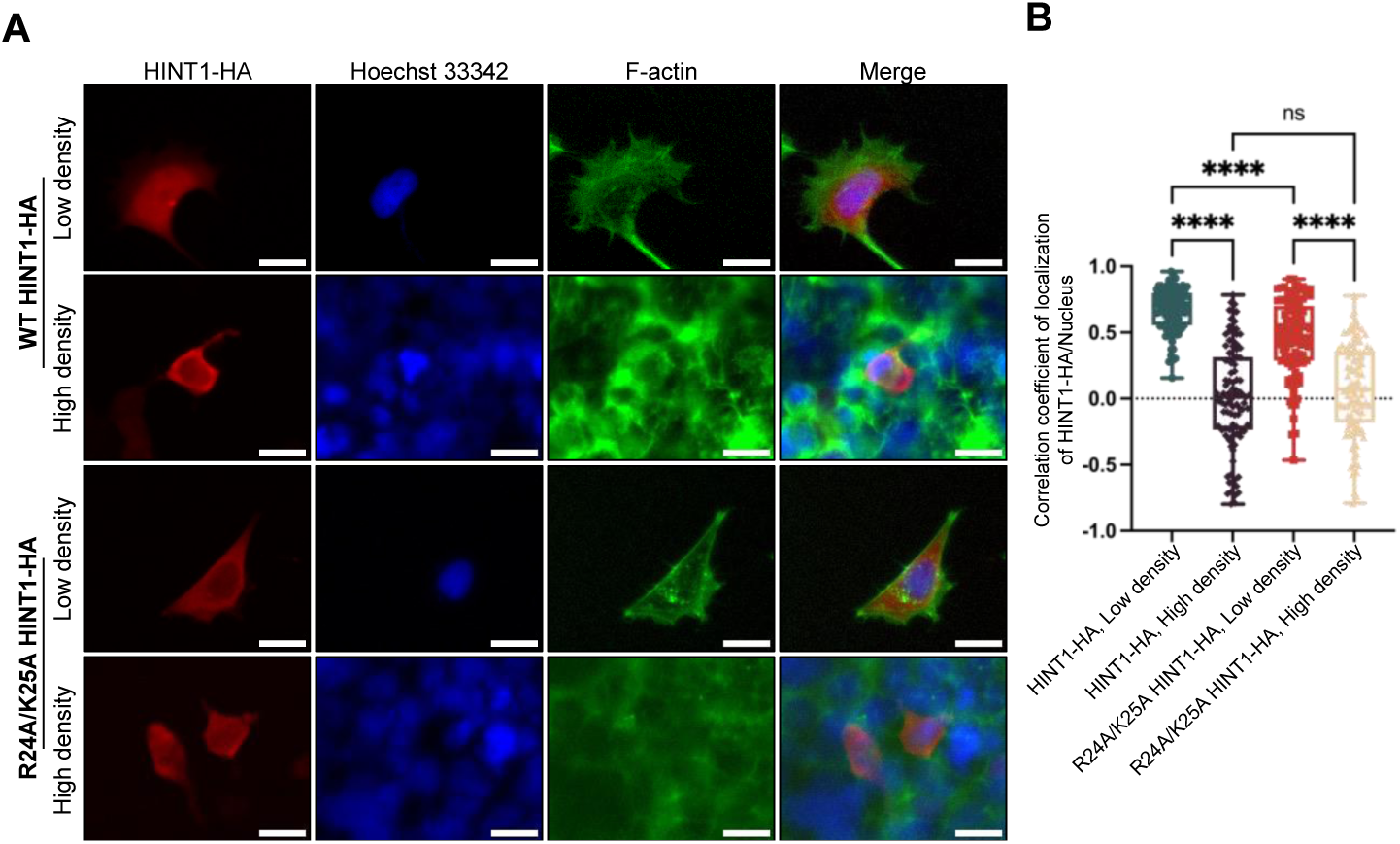
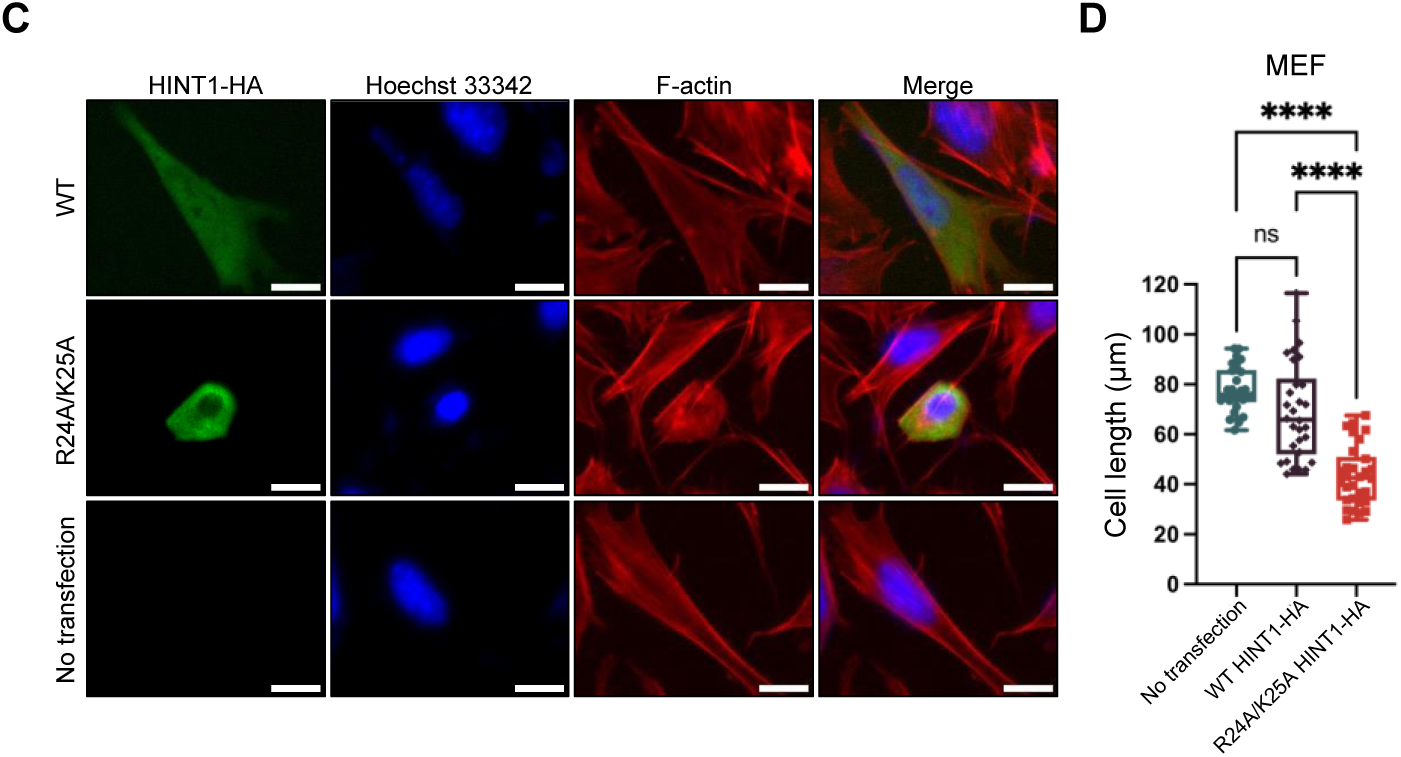
Forced expression of HINT1 in the cytoplasm disrupts elongation of cells (**A** and **B**) Exogenously expressed HINT1-HA in HINT1-KO HEK293A cells at low density primarily localizes in the nucleus, whereas the point mutation in the intramolecular NLS (R24A/K25A) promotes its cytoplasmic localization. The cells were stained as shown in Fig. 1A, and the correlation coefficient between HINT1-HA staining and Hoechst nuclear staining was calculated and plotted. Scale: 20 μm. (**C** and **D**) MEF cells were transfected with WT and mutant R24A/K25A HINT1-HA at low density and stained with rabbit anti-HA antibody followed by anti-rabbit IgG- Alexa Fluor Plus 488 antibody (green). Nuclear DNA was stained with Hoechst 33342 (blue). Actin filaments were stained with phalloidin Alexa Fluor 568 (red) Notably, the mutant HINT1 localized in the cytoplasm and disrupted the elongated shape of the cells, while the WT HINT1 did not affect cell morphology. The length of the major axis of each cell was measured and plotted. Each dot represents the length of an individual cell (n=30). Scale: 20 μm.

### Transloction of HINT1 to the cytoplasm is regulated by exportin 1 downstream of high cell density

Point mutation of intramolecular NLS of HINT1 disrupted its nuclear translocation, suggesting that active transport is involved in the nucleocytoplasmic shuttling of HINT1. First, we tested two available importin inhibitors—ivermectin (an importin α/β1 complex inhibitor) and importazole (an importin β inhibitor) [26]. However, neither showed a significant effect on the localization of HINT1 (**Fig. S8**). Next, the cells were treated with Leptomycin B, an exportin 1/CRM1 inhibitor, at both low and high cell densities. Leptomycin B

did not affect the nuclear localization of HINT1 at low density but inhibited its translocation to the cytoplasm at high density (**Fig. 6A and 6B**). At high cell density, the inhibition of exportin 1 led to an increase in cell cross-sectional area in WT cells, but this effect was not observed in HINT1 KO cells (**Fig. 6C and 6D**).

**Fig. 6.**
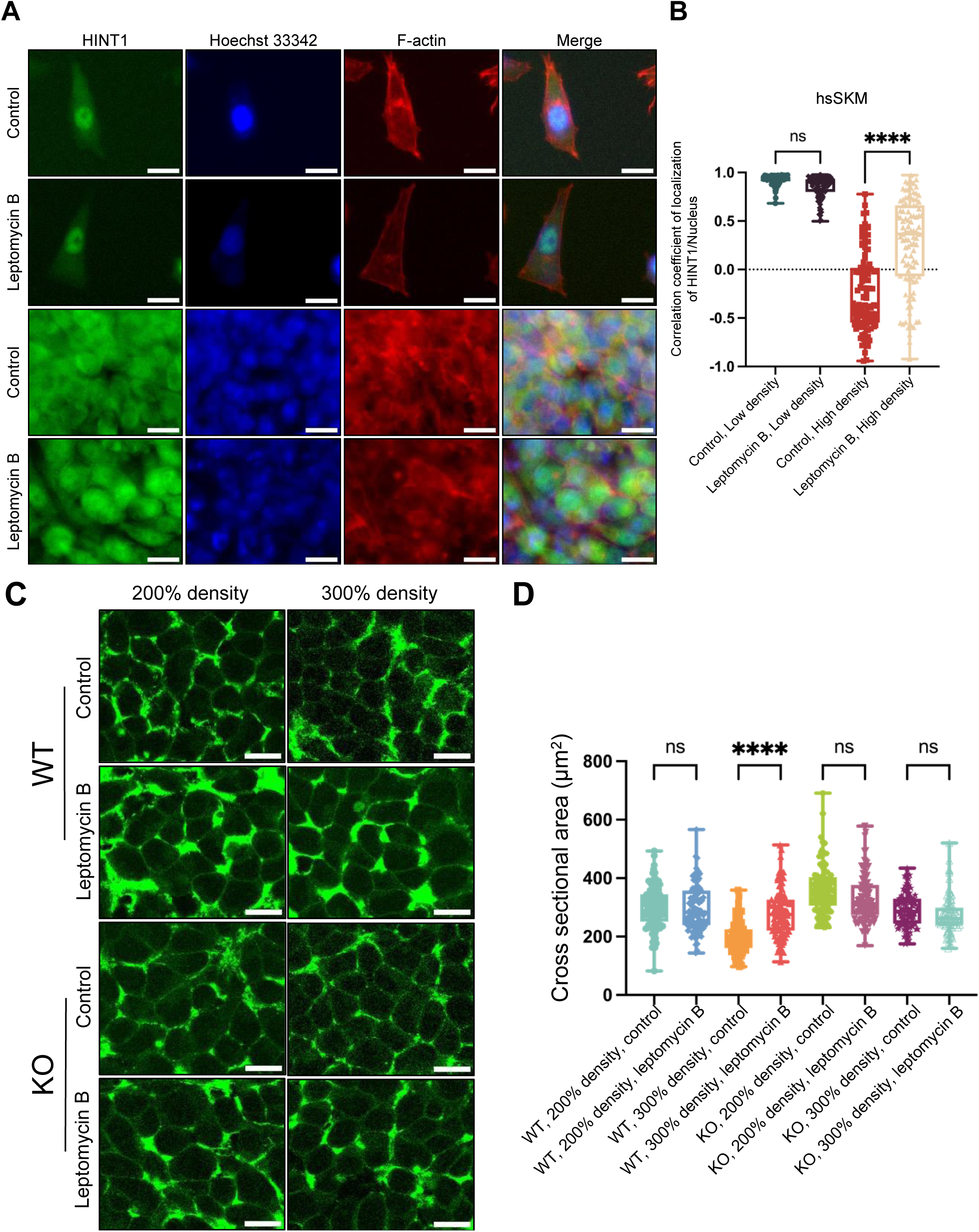
The inhibition of exportin 1 by Leptomycin B blocks the translocation of HINT1 to the cytoplasm at high cell density and increases the cell cross-sectional area (A) hsSKM cells were treated with or without 10 nM of Leptomycin B for 24 hours at both low and high densities, then stained with rabbit anti-HINT1 antibody followed by anti-rabbit IgG- Alexa Fluor Plus 488 antibody. Nuclear DNA was stained with Hoechst 33342. Actin filaments were stained with phalloidin Alexa Fluor 568.. Scale: 20 μm. (B) Correlation coefficient of HINT1 staining and Hoechst nuclear staining were calculated and plotted. Similar results were obtained with HEK293A and MEF cells (Fig. S9). (C) WT and HINT1-KO HEK293A cells were treated with or without 10 nM of Leptomycin B for 24 hours at high densities and stained with fluorogenic membrane dye to demarcate the cell boundaries. Scale: 20 μm. Values represent the cross-sectional area of each cell as a dot (n=100). The percentages are values used for convenience when seeding cells and do not represent the actual cell density. However, for 100%, the cell count was measured and cultured in such a way that the cells would reach confluency by the time of adding Leptomycin B.

### Translocation of HINT1 to the cytoplasm inhibits PKC

Although increased PKC activity has been observed in HINT1-deficient mice [27], the role of HINT1 translocation in regulating PKC activity has not yet been explored. We hypothesized that PKC activity is elevated at low cell density due to HINT1’s localization in the nucleus, while at high density, PKC activity decreases as translocated HINT1 inhibits PKC in the cytoplasm. Since myristoylated alanine-rich C kinase substrate (MARCKS) is a major PKC substrate, with PKC-dependent phosphorylation of MARCKS playing a key role in cytoplasmic actin remodeling [28, 29], we first measured the phosphorylation levels of MARCKS in various cell lines at low and high densities using a phospho-MARCKS (Ser167/170) polyclonal antibody [30]. As anticipated, the phosphorylation levels of MARCKS were significantly higher in low-density cells compared to high-density cells (**Fig. 7A**). Next, we tested whether forced expression of HINT1 in the cytoplasm of low-density cells would also inhibit MARCKS phosphorylation. Compared to non-transfected cells, overexpression of WT HINT1-HA led to a decrease in MARCKS phosphorylation, likely due to the presence of some HINT1 in the cytoplasm (**Fig. 7B**). Supporting our hypothesis, the mutant form of HINT1, which predominantly localizes in the cytoplasm, resulted in an even greater reduction in MARCKS phosphorylation.

**Fig. 7.**
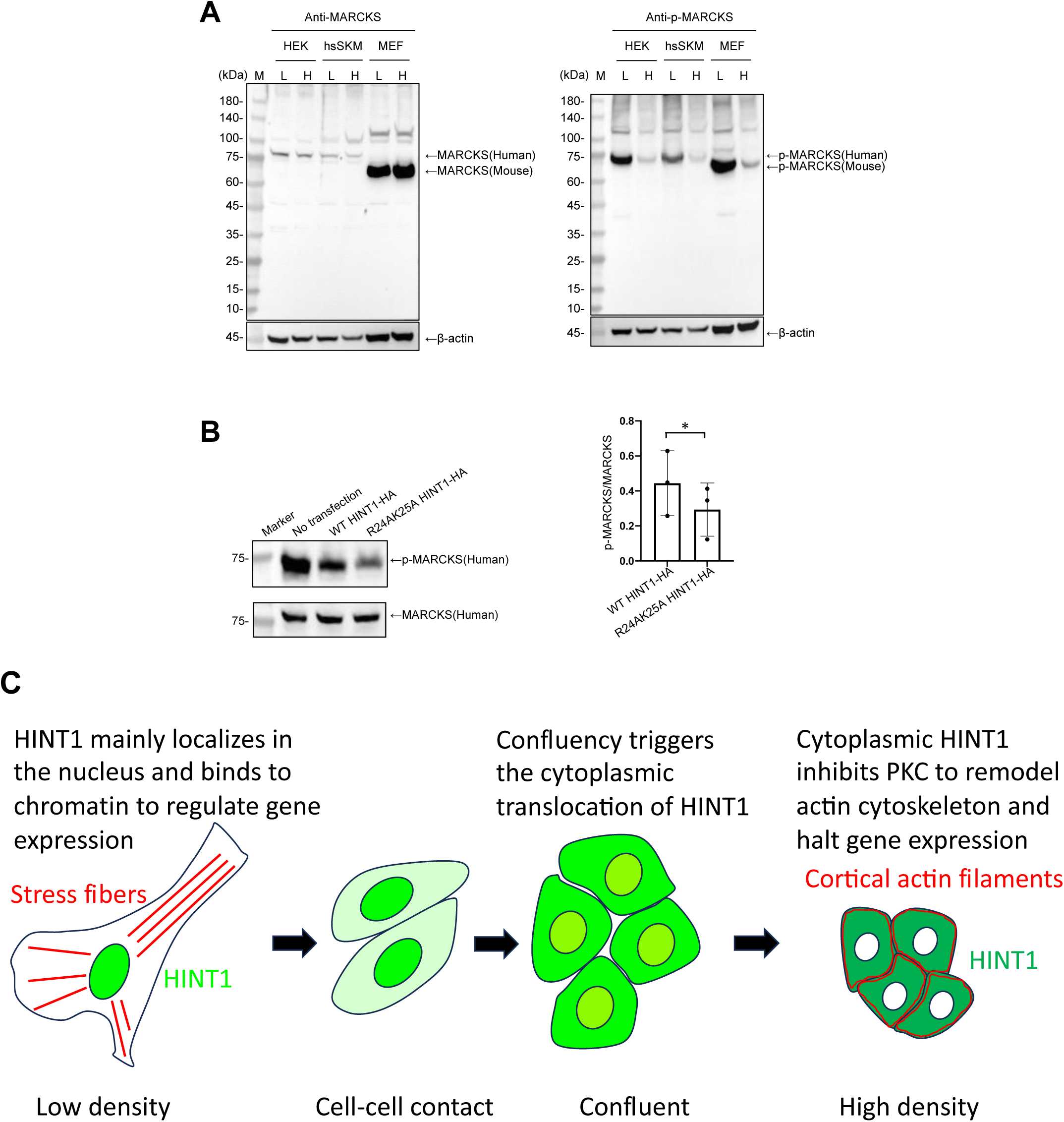
Translocation of HINT1 to the cytoplasm inhibits PKC activity (A) MARCKS phosphorylation is dependent on cell density in HEK293A, hsSKM, and MEF cells. Cells were cultured overnight at low and high densities, and the phosphorylation levels of MARCKS were assessed by western blotting using a phospho-MARCKS (Ser167/170) polyclonal antibody. Total MARCKS and actin levels were also measured via western blotting as internal controls. (B) Cytoplasmic forced-expression of HINT1 reduces MARCKS phosphorylation. HEK293A cells were seeded at low density, transfected with either WT HINT1-HA or mutant R24A/K25A HINT1-HA, incubated for 24 hours, and subsequently lysed in SDS sample buffer. Phospho-MARCKS, total MARCKS, and actin levels were analyzed via western blotting. Quantitative analysis of MARCKS phosphorylation levels in HEK293 cells, with or without transfection of WT HINT1-HA or mutant R24A/K25A HINT1-HA. The data was normalized using the band intensity from non-transfected cells. n = 3. Statistical analysis was conducted using paired t-test. * P < 0.05. (C) A proposed model illustrates how HINT1 translocation is regulated by cell density and how this process contributes to the disruption of actin SFs, leading to a rounded, confined cell morphology. At low density, HINT1 predominantly localizes in the nucleus, where it binds to chromatin to regulate gene expression, thereby promoting cell proliferation and development. Cell-cell contact alone is insufficient to induce HINT1 translocation to the cytoplasm. At confluency, cells initiate the export of HINT1 to the cytoplasm, a process mediated by exportin-1. As cell density increases, more HINT1 is actively transported to the cytoplasm. In the cytoplasm, HINT1 inhibits PKC, which influences actin remodeling through various signaling pathways, resulting in a rounded and compact cell morphology. This translocation also suppresses gene expression, leading to a halt in proliferation and promoting CIP. The subcellular localization of HINT1 appears to be cell type-dependent, which may explain why some cells form SFs while others do not *in situ*.

## Discussion

We recently developed a DSP-MNase proteogenomics method to identify chromatin-binding proteins that respond to mechanical stress and cell density [2, 19]. Among the proteins identified was HINT1 from sub- confluent (∼80% confluency) hsSKM cells, although its role in mechanotransduction had not been previously explored. While most proteins known to shuttle between the nucleus and cytoplasm in a cell density-dependent manner are also sensitive to mechanical force and substrate stiffness [2], we found that HINT1 is uniquely responsive to cell density but not to mechanical force.

HINT1 was originally identified as a tumor suppressor and an inhibitor of PKC [16]. HINT1 is a member of the histidine triad (HIT) family, proteins that are recognized for their roles in nucleotide binding and hydrolysis [31]. HINT1 also acts as a transcriptional suppressor via direct binding to transcription factors [32–34].

HINT1 was first noted for its involvement in regulating cell growth and signal transduction and has since been linked to several key cellular functions, including apoptosis, gene expression regulation, and cellular stress response mechanisms [17]. Although HINT1 KO mice exhibit normal fetal and adult development, they show increased susceptibility to tumor formation and display increased anxiety-like behavior along with impaired motor coordination [35–38].

HA-tagged HINT1 was initially reported to localize primarily in the cytoplasm of a human fibroblast cell line, with minimal presence in the nucleus [39]. However, a subsequent study found that endogenous HINT1 predominantly localizes in the nucleus [40]. Our findings may provide an explanation for this discrepancy, as transfected cells often reach high densities, even though only the transfected cells are visible via fluorescent probes. Additionally, while HINT1 was primarily detected in the nucleus in human breast tissue sections, suggesting it does not function as a PKC inhibitor [40], this nuclear localization appears to occur in actively growing epithelial cells but not in deeper quiescent cells.

Given its small size (147 amino acid residues), it was surprising to find that HINT1 is not a mechanosensitive nucleocytoplasmic shuttling protein that can be imported through the extended nuclear pore complex (NPC). Although HINT1 forms dimers [41], its overall molecular weight still appears small enough to pass through NPC. According to the NPC stretching model [42], small molecules could be passively transported through a mechanically stretched NPC. For instance, we recently identified ubiquitin-conjugating enzyme E2 A (152 residues) and core-binding factor subunit beta (182 residues) as mechano- and cell density-sensitive nucleocytoplasmic shuttling proteins [19, 22, 43]. In contrast, HINT1 translocation appears to be actively regulated by a transporter [26]. Pharmacological experiments showed that HINT1’s nuclear export is governed by exportin 1/CRM1, which is activated by high cell density but not by cell-cell contact, through an unknown pathway. Although the importin responsible for HINT1’s nuclear import remains unidentified, importin α/β1 has been ruled out based on pharmacological testing. A point mutation in the predicted intramolecular NLS impaired HINT1’s nuclear entry at low density, indicating that an unknown importin must be involved in this process.

We also discovered that at high cell density, HINT1 translocation to the cytoplasm triggers the collapse of actin SFs, leading to a morphological shift in the cells, resulting in a smaller, rounder shape. This is also confirmed by forced expression of HINT1 in the nucleus at low density, whereas normal HINT1 is mainly localized in the nucleus. Retaining HINT1 in the nucleus may not only promote its function for gene expression but also promote PKC activity in the cytoplasm. Active PKC in the cytoplasm may induce actin SFs through multiple pathways because many actin-binding proteins and signaling molecules involved in actin remodeling are PKC substrates [44, 45]. For example, in addition to the phosphorylation-dependent remodeling of actin filaments mediated by MARCKS, the PKC-AKT-Girdin pathway may also promote SF formation, as Girdin phosphorylation enhances SF assembly [46]. Furthermore, HINT1 knockdown has been shown to increase PKC activity, as well as elevate both the expression and phosphorylation levels of AKT and Girdin [27, 47].

Translocation of HINT1 to the cytoplasm at high density may inhibit the PKC-mediated SF formation because HINT1 binds to a full-length PKC and inhibits its kinase activity [40, 48]. Interestingly, reduction of HINT1 expression is observed in cardiac hypertrophic patients and the cross-sectional area of cardiomyocytes from HINT1-KO mice was larger than that of control mice in response to hypertrophic stimuli [48], which is consistent with our observation in HINT1 KO cells.

In conclusion, we have discovered a novel regulatory mechanism within the long-established PKC signaling pathway. Specifically, our findings demonstrate that nucleocytoplasmic translocation of HINT1 regulates the disassembly of SF, through the PKC-MARCKS axis (**Fig. 7C**). At low cell density, HINT1 remains in the nucleus, where it does not inhibit PKC in the cytoplasm and binds to chromatin to regulate gene expression. Although cell-cell contact alone is insufficient to trigger HINT1’s translocation, near confluency induces its movement to the cytoplasm. As cell density increases, more HINT1 translocates to the cytoplasm, leading to a reduction in PKC activity. This decreased PKC activity remodels actin filaments, at least partially through MARCKS dephosphorylation, resulting in a more rounded and compact cell shape that supports the maturation of tissue formation. Simultaneously, the removal of HINT1 from the nucleus suppresses gene expression, thereby promoting CIP. These findings are consistent with previous observations that CIP in Madin-Darby canine kidney cells proceeds in three distinct phases [3], yet our results further suggest that the final phase involves HINT1-driven actin remodeling at high cell density, independent of mechanical constraints.

Our work paves the way for future investigations into the molecular mechanisms by which HINT1 inhibits specific PKC isoforms, the role of cell density in activating exportin and importin, distinctions between cell-cell contact, confluency, and high density regarding intracellular structure and biological function, the factors driving these transitions, and how HINT1 loss prolongs tissue culture cell survival under high-density conditions.

## Materials and methods

### Antibodies and reagents

Rabbit monoclonal anti-HINT1 antibody was purchased from Abcam (AB124912). Mouse monoclonal anti-YAP/TAZ antibody was purchased from Santa Cruz Biotechnology (sc-101199). Rabbit polyclonal anti-MARCKS (10004-2-Ig) and phospho (Ser167/170)-MARCKS (29145-1-AP) were purchased from Proteintech. Anti-Rabbit IgG- Alexa Fluor Plus 594 antibody (A32754), Anti-Rabbit IgG- Alexa Fluor 488 antibody (A11034), Anti-Mouse IgG- Alexa Fluor Plus 594 antibody (A32744), Phalloidin Alexa Fluor 488 (A12379), and Phalloidin Alexa Fluor 568 (A12380) were purchased from Invitrogen. Mouse monoclonal anti-β-actin antibody (HC201-01), Goat Anti-Rabbit IgG-HRP antibody (HS101-01), and Goat Anti-Mouse IgG-HRP antibody (HS201-01) were purchased from Transgen. Hoechst 33342 was purchased from Thermo Fisher Scientific. Cellbrite Fix 488 membrane stain was purchased from Biotium (30090-T). Latrunculin B was purchased from Sigma-Aldrich (428020). Blebbistatin was purchased from AbMole BioScience (856925-71-8). Leptomycin B was purchased from Solarbio (87081-35-4). Importazole (HY-101091) and Ivermectin (HY- 15310) were purchased from MedChemExpress.

### Plasmid construction

Human HINT1 cDNA (UniProt Accession ID: P49773) was amplified by PCR using 5’ primer, 5′- CCCAAGCTTATGGCAGATGAGATTGCCAAGGC-3′, and 3’ primer, 5′- CCGGAATTCACCAGGAGGCCAATGCATTTG-3′, using HEK293A cDNA library as a template and ligated into pcDNA3-HA vector at HindIII/EcoRI sites [49].

NLS-HINT1 and mutant NLS-HINT1 were constructed by PCR using 5’ primer, 5′- CCCAAGCTTATGCCAAAAAAGAAGAGAAAGGTAATGGCAGATGAGATTGCCAAGGC-3′, 3’ primer, 5′-CCGGAATTCTTAACCAGGAGGCCAATGCATTTG-3′, and 5’ primer, 5′- CCCAAGCTTATGCCAAAAACGAAGAGAAAGGTAATGGCAGATGAGATTGCCAAGGC-3′, 3’ primer, 5′-CCGGAATTCTTAACCAGGAGGCCAATGCATTTG-3′, respectively and ligated into pcDNA3-HA vector at HindIII/EcoRI sites.

Similarly, R24A/K25A mutant HINT1-HA was constructed using 5’ primer, 5′- CGATCTTTGGGAAGATCATCGCGGCGGAAATACCAGCCAAAATCATTTTTG-3′, and 3’ primer, 5′- GATGATCTTCCCAAAGATCG-3′, and ligated into pcDNA3-HA by T5 exonuclease DNA assembly (TEDA) method [50].

### Cell culture and transfection

HEK293A cells were purchased from Thermo Fisher. Human skeletal muscle (hsSKM) and mouse embryonic Fibroblast (MEF) cells were purchased from ATCC. These cells were grown in DMEM (Biological industries, Israel) supplemented with 10% FBS (Biological industries, Israel) and 1% penicillin-streptomycin. Cells were maintained at 37°C and 5% CO_2_. Cells were transfected with polyethylenimine (PEI) or LipoGene2000 Star transfection reagent (L7002, US Everbright) in accordance with the manufacturer’s protocol.

### Immunofluorescence microscopy

Cells were plated on a 48-well plate coated with or without 10 ng/ml fibronectin (03-090-1-01, Biological Industries, Israel), washed once with PBS-D (PBS containing 1 mM Ca²⁺ and Mg²⁺), fixed with 4% formaldehyde in PBS-D for 20 minutes, rinsed in PBS-D, permeabilized with 0.5% Triton X-100 in TBS (50 mM Tris-HCl, pH 7.4, 150 mM NaCl) for 10 minutes, then rinsed again in TBS-0.1%Tx (TBS containing 0.1% Triton X-100), and blocked with 2% BSA in TBS-0.1%Tx for 1 hour. Cells were then incubated with primary antibodies for 2 hours, followed by several washes with TBS-0.1%Tx, incubated with secondary antibodies, and washed several more times with TBS-0.1%Tx. Cells were further incubated with Phalloidin Alexa Fluor 488 or Phalloidin Alexa Fluor 568, washed multiple times with TBS-0.1%Tx, and stained with Hoechst 33342 for 15 minutes. After a final wash, cell imaging was performed using the EVOS® FL Auto Imaging System (Thermo Fisher) with an EVOS® Obj, Inf Plan Fluor 20 LWD objective, and Plan Fluor 40 LWD, .65NA/2.8WD objectives. Image processing and analysis were conducted using NIH Image J software (version 1.54g).

### Immunofluorescence confocal microscopy

WT and HINT1-KO HEK293A cells were cultured on cover glasses coated with either polylysine or fibronectin. The cells were stained with Cellbrite Fix 488 membrane stain for 15 minutes at 37°C, then fixed as described above and prepared for imaging by being mounted with mounting medium (Spring Bioscience). Imaging was performed using a Leica SP8 confocal microscope equipped with a 63X HC PL APO objective lens with a numerical aperture (NA) of 1.40. The acquired images were processed and analyzed using Image J software.

### Western blotting

Cells were lysed in SDS or LDS sample buffer and heated at 95°C for 5 minutes. The samples were loaded onto either a homemade SDS-PAGE gel or a Novex NuPAGE 4–12% gradient Bis-Tris gel. Separated proteins were transferred to a nitrocellulose membrane and blocked with blocking buffer (5% non-fat milk powder in TBST: 20 mM Tris–HCl, pH 7.4, 110 mM NaCl, 5 mM MgCl_2_, 0.1% Tween 20) for 2 hour. Primary antibodies were prepared in this blocking solution, and the membranes were incubated overnight at 4°C. After washing in TBST, the membranes were incubated with HRP-conjugated secondary antibodies in blocking solution for 1 hour at room temperature. The membranes were then washed and developed using HRP substrate (WesternBright ECL, Advansta).

### Knock-out of HINT1 in tissue culture cells

HINT1 KO HEK293A cells were generated by delivery of Cas9 and target-specific guide RNAs (gRNAs). Oligos encoding the gRNAs for HINT1 were designed using CRISPick (https://portals.broadinstitute.org/gpp/public/analysis-tools/sgrna-design) and the selected HINT1-specific gRNAs sequence, 5′-CACCGCCTGGTGGCGACACGATCTT-3′ and 5′- AAACAAGATCGTGTCGCCACCAGGC-3′, were cloned into BbsI digested pX330-U6-Chimeric_BB-CBh- hSpCas9 (Addgene plasmid ID: 42,230). Px330-sgHINT1-exon1 plasmids were transfected into HEK293A cells using LipoGene2000 Star transfection reagent (US Everbright) according to manufactures’s protocol.

Briefly, HEK293A cells were seeded into a 24-well plate. After 24 h that the cells reached 60%–70% confluent, 0.8 μg Px330-sgHINT1-exon1 plasmid was added to the well in the presence of LipoGene2000 Star transfection reagent. 72h post-transfection, cells were then separated as single cells into a 96-well plate by serial dilution for another 7 days. Individual clones were expanded, and HINT1 protein expression was examined by immunoblotting.

### RNA-sequencing

WT and HINT1 KO HEK293A cells were seeded at 5×10^5^ cells in 60 mm dishes and cultured in DMEM supplemented with 10% FBS and 1% Pen/Strep for 18–24 h. Total RNA was isolated using the RNAisoPlus (Takara) and sent to Beijing Biomarker Technologies Co., Ltd. for library construction and sequencing. Briefly, extracted high quality RNA were used as input material for the RNA sample preparations. Sequencing libraries were generated according to the following steps. Firstly, mRNA was purified from total RNA using poly-T oligo-attached magnetic beads. Fragmentation was carried out using divalent cations under elevated temperature in a fragmentation buffer. First strand cDNA was synthesized using random oligonucleotides. Second strand cDNA synthesis was subsequently performed using DNA Polymerase I and RNase H. Remaining overhangs were converted into blunt ends via exonuclease/polymerase activities and the enzymes were removed. After adenylation of the 30 ends of the DNA fragments, adapter oligonucleotides were ligated to prepare for hybridization. After post-Ligation purification, DNA fragments with ligated adaptor molecules on both ends were selectively enriched in a 15 cycle PCR reaction. Then, after post amplification purification and quality control, the sequencing library was sequenced on Illumina NovaSeq 6000 platform by Beijing BioMarker Technologies Cp. Ltd.

### Cell migration assay

For random cell migration, WT and HINT1 KO HEK293A calls were seeded at 1 ×10^5^ cells per well on a 6- well plate. After 24 hours, images were acquired for 20 h at 1 frame/10 min at 37°C using a ×10 Plan FL objective on an EVOS® FL Auto time lapse microscope with a monochrome and color camera. Cells were tracked using ImageJ (Plugin: Manual tracking) to obtain migration speed (μm/min). Cells that died, divided, or moved out of the frame were excluded from the analysis and tracking. The path of each cell was obtained as a track using ImageJ (Plugin: Chemotaxis tool).

### Quantification of intracellular protein distribution

To analyze the translocation of HINT1, the intensity of fluorescence along a predetermined line in an immunofluorescent image, which shows the nucleus in blue and the protein of interest in red, was measured using the “Plot Profile” function in ImageJ. To assess the relationship between the intensity levels in the blue (nucleus) and red (protein of interest) channels, and thereby gain insights into the localization of the protein within the cell, the Pearson’s correlation coefficient was calculated. This coefficient ranges from -1, indicating a complete cytosolic distribution, to 1, which signifies an exclusive nuclear presence, using GraphPad Prism

10.1.1. To analyze cross sectional area, the demarcation of cell boundaries were completed by Cellpose 3 software (https://www.cellpose.org) [51]. Then, the cells were randomly selected and measured area using the measurement function of ImageJ.

## Data analysis

For RNA-seq analysis on Galaxy (https://usegalaxy.org), raw reads were trimmed using TrimGalore (Galaxy Version 0.6.7+galaxy0) and quantified transcript abundance using Salmon quant (Galaxy Version 1.10.1+galaxy2). Summarized transcript-level data were performed using tximport function (Galaxy Version 1.22.0). Differential expression and enrichment analysis were performed on iDEP (http://bioinformatics.sdstate.edu/idep/).

## Statistics

Data are mean ± S.D. All experiments were performed at least three times independently. All image analysis was performed by operators who were blinded to the treatments administered. Significance was analyzed by t-test, one-way ANOVA followed by Tukey’s multiple comparison test, two-way ANOVA followed by Tukey’s multiple comparison test, or three-way ANOVA followed by Tukey’s multiple comparison test. * P≦0.05, ** P ≦0.01, *** P≦0.001, **** P≦0.0001, ns: not significant. Statistical analysis was performed in graphPad Prism 10.1.1.

## Acknowledgments

## Acknowledgements

We thank a support of Tianjin university SPST core facility.

## Author contributions

Conceptualization: F.N.; sample preparation and data acquisition: X.Z.; analysis: X.Z.; writing: X.Z. and F.N.

## Data and materials availability

Data that support the findings of this study are available within the article and its Supplementary figures and tables. The RNA-seq data generated from this study have been deposited in the Gene Expression Omnibus (GSE270952, temporary reviewer login: https://www.ncbi.nlm.nih.gov/geo/query/acc.cgi?acc=GSE270956, secure token: qlwbysgwrxsxpqj). All other data supporting the findings of this study are available from the corresponding author on reasonable request. Materials may be requested from the corresponding author.

**Supplemental material** Supplementary Fig. S1–S9. Table S1: Primers used.

Table S2: Excel file containing additional data too large to fit in a PDF, related to Fig. 3. Source Data for Fig. 1c and 7b.

---

**Fig. S1.**
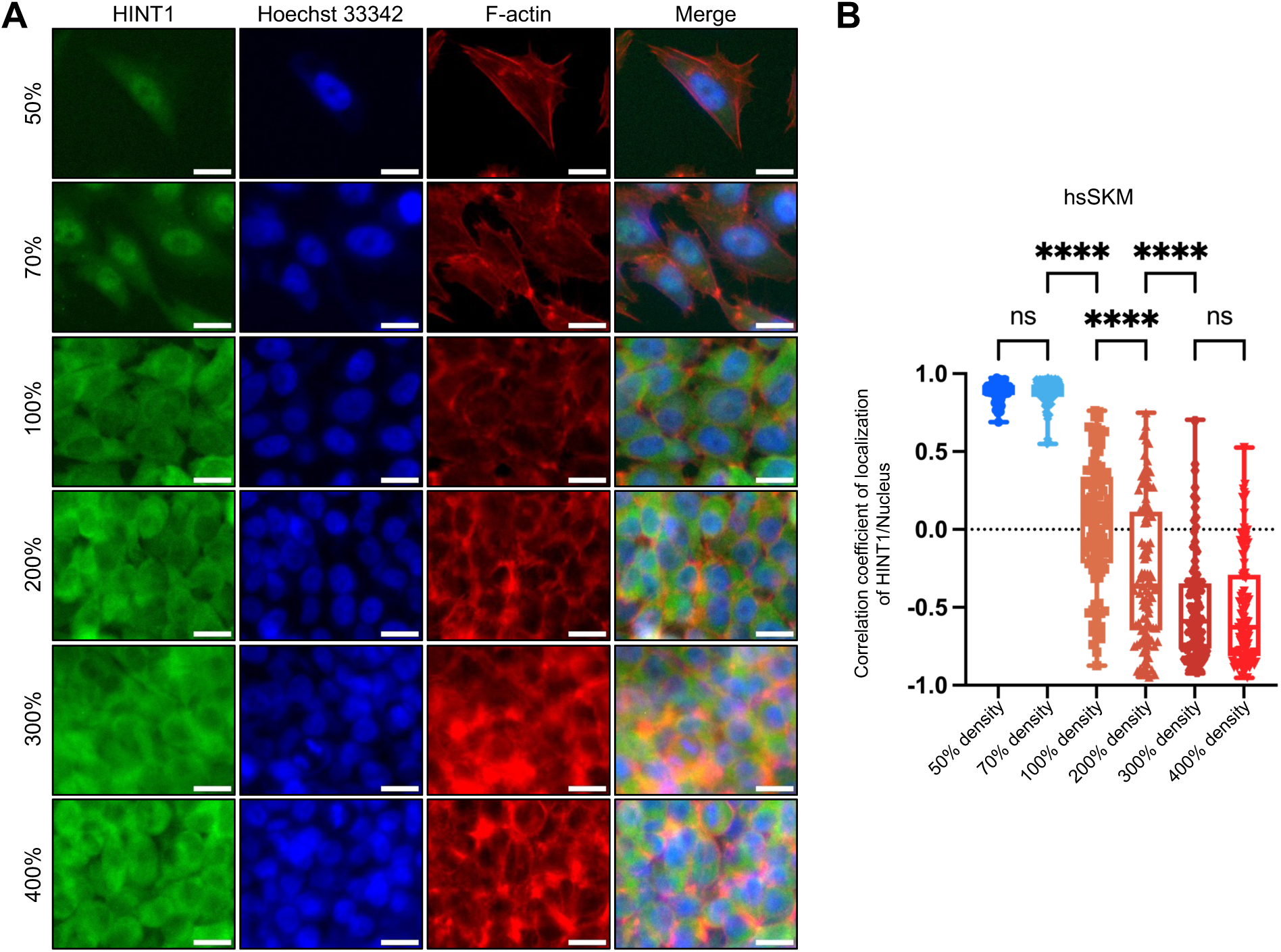
The translocation of HINT1 is not dependent on cell-cell contact but requires a high cell density (A) hsSKM cells were cultured at different density. After fixation, HINT1 was stained by anti- HINT1 antibody followed Alexa-488 secondary antibody. The nucleus was stained with Hoechst 33342 and F-actin was stained with phalloidin conjugated with 568. Scale: 20μm. (B) Correlation coefficient of HINT1 staining and Hoechst 33342 staining were calculated and plotted. Note that HINT1 remains in the nucleus when cells are in partial contact with each other (e.g., 70% confluency). However, near full confluency (e.g., 100%, with some space still visible between cells) is sufficient to trigger its translocation to the cytoplasm. As cell density increases, more HINT1 moves to the cytoplasm. The percentages are values used for convenience when seeding cells and do not represent the actual cell density. However, for 100%, the cell count was measured and cultured in such a way that the cells would reach confluency by the time of fixation.

**Fig. S2.**
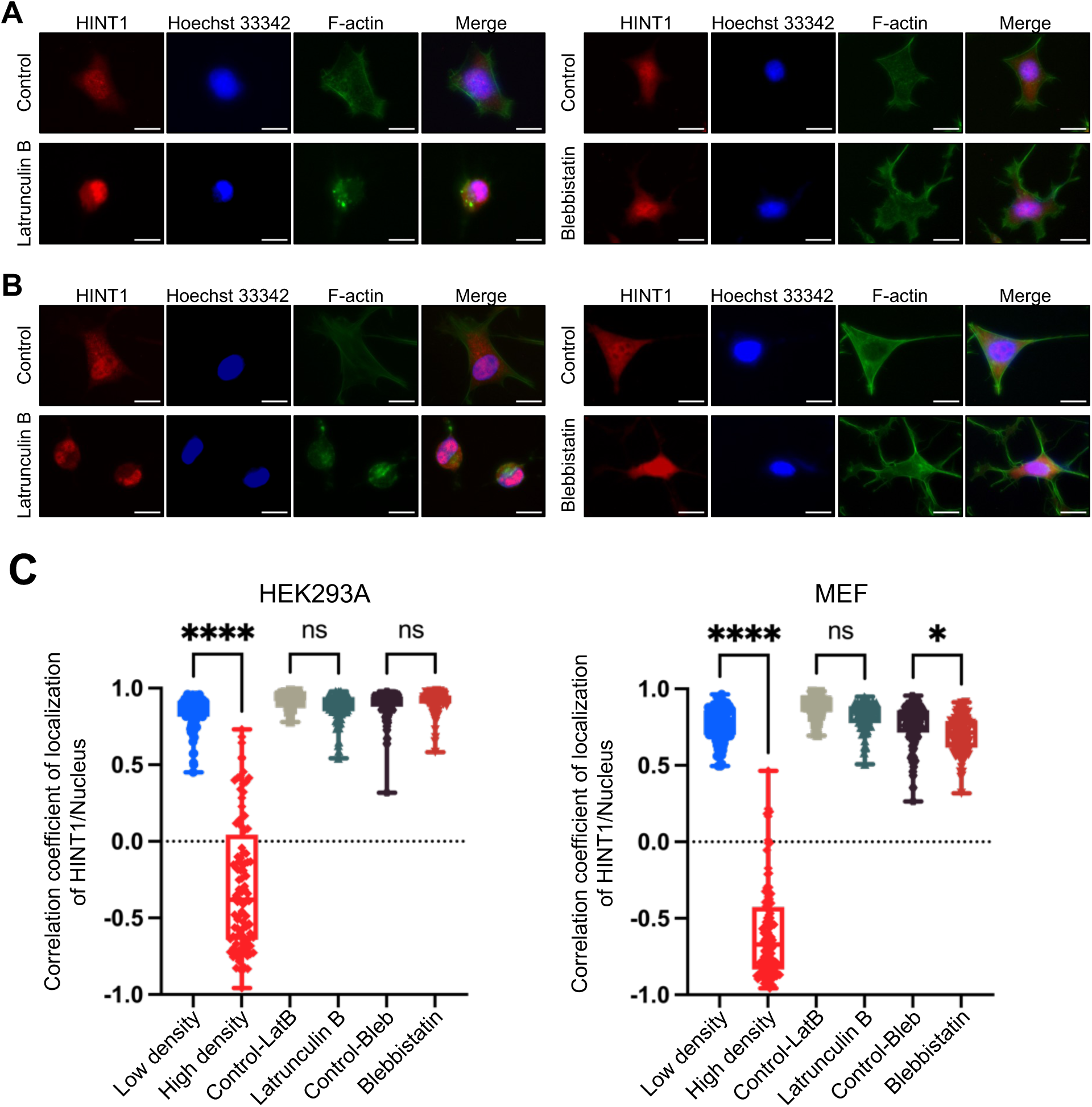
Nucleocytoplasmic translation of HINT1 is not mechanosensitive (A) HEK293A cells were treated with or without 5 μM of Latrunculin B for 60 min or 50 μM blebbistatin for 180 min. Scale: 20 μm. The cells were stained as shown in Fig. 1D. Correlation coefficient of HINT1 staining and Hoechst 33342 staining were calculated and plotted. (B) MEF cells were treated with or without 5 μM of Latrunculin B for 60 min or 50 μM blebbistatin for 180 min. Scale: 20 μm. The cells were stained as shown in Fig. 1D. (C) Correlation coefficient of HINT1 staining and Hoechst 33342 staining were calculated and plotted.

**Fig. S3.**
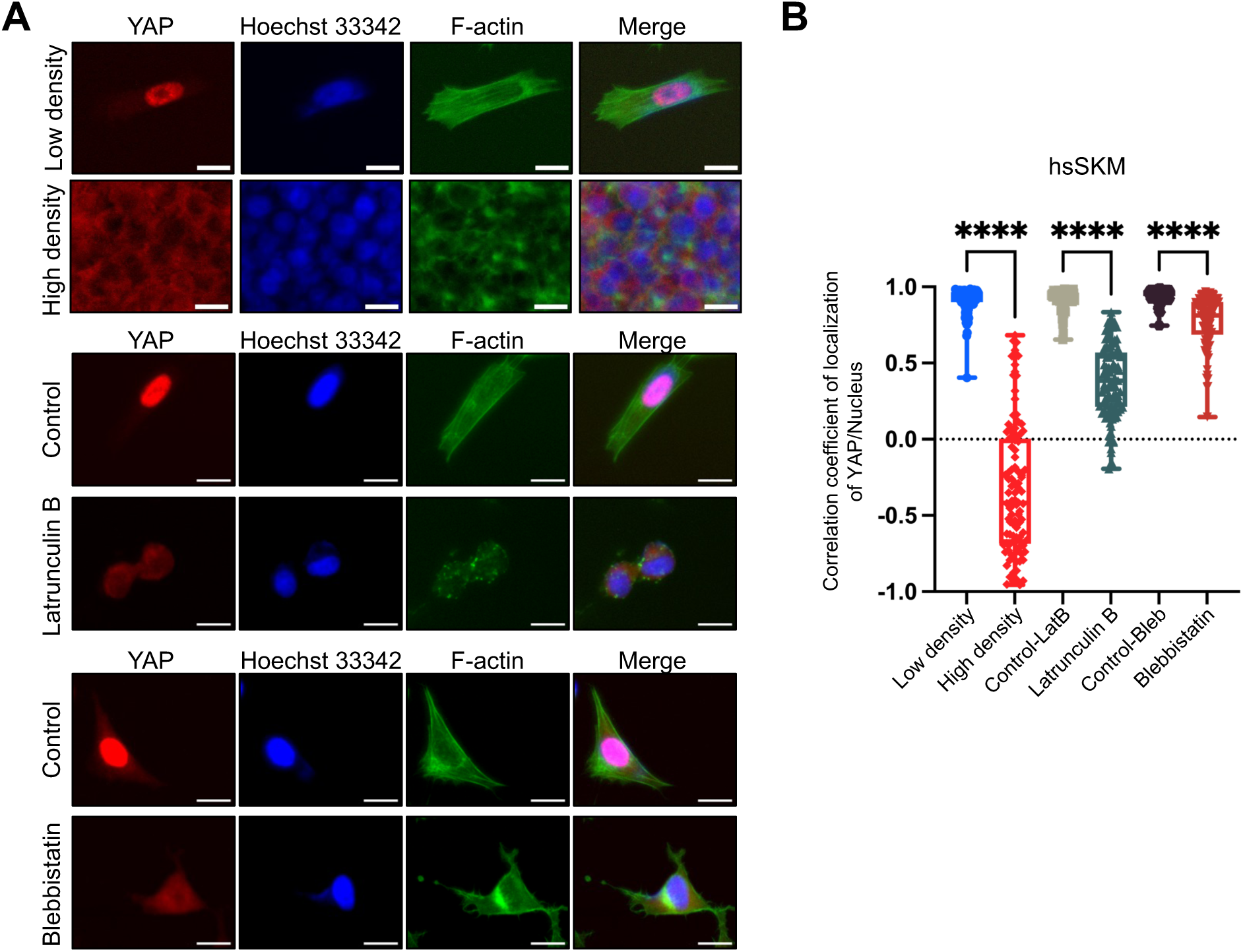
Nucleocytoplasmic translocation of YAP is sensitive to cell density, F-actin and myosin contraction (A) hsSKM cells were cultured at low or high density (top two panels). hsSKM cells were treated with or without 5 μM of Latrunculin B for 60 min (middle two panels). The cells were treated with or without 50 μM of blebbistatin for 180 min (bottom two panels). Scale: 20μm. YAP was stained with mouse anti-YAP antibody followed by anti-mouse IgG- Alexa Fluor Plus 594 antibody (red). Nuclear DNA was stained with Hoechst 33342 (blue). Actin filaments were stained with phalloidin Alexa Fluor 488 (green). (B) Correlation coefficient of YAP staining and Hoechst 33342 staining were calculated and plotted.

**Fig. S4.**
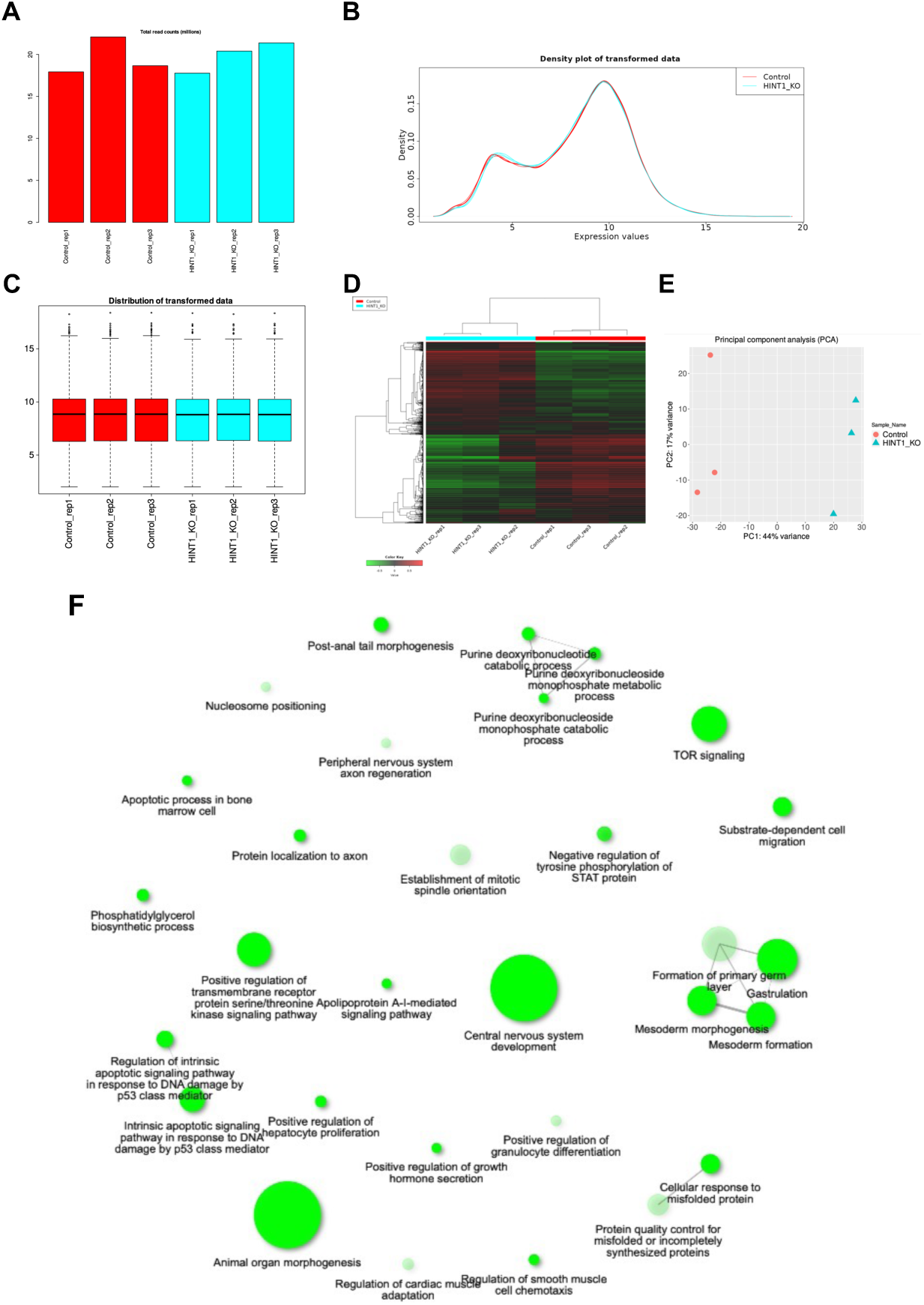
RNAseq analysis of HINT1 KO HEK293A cells Diagnostic plots for read-counts data. (A) A bar plot of total read counts per library, showing some variation in library sizes. (B) Distribution of transformed data using a density plot. (C) Boxplot of transformed data. Variation among replicates is small (B and C). (D) Hierarchical clustering. (E) Principal component analysis (PCA) analysis indicate that there are clear differences in gene expression between WT and HINT1-KO cells. (F) The interactive plot showing the relationship between enriched down-regulated pathways in HINT1-KO vs WT cells. Two pathways (nodes) are connected if they share 30% (default, adjustable) or more genes. Darker nodes are more significantly enriched gene sets. Bigger nodes represent larger gene sets.

**Fig. S5.**
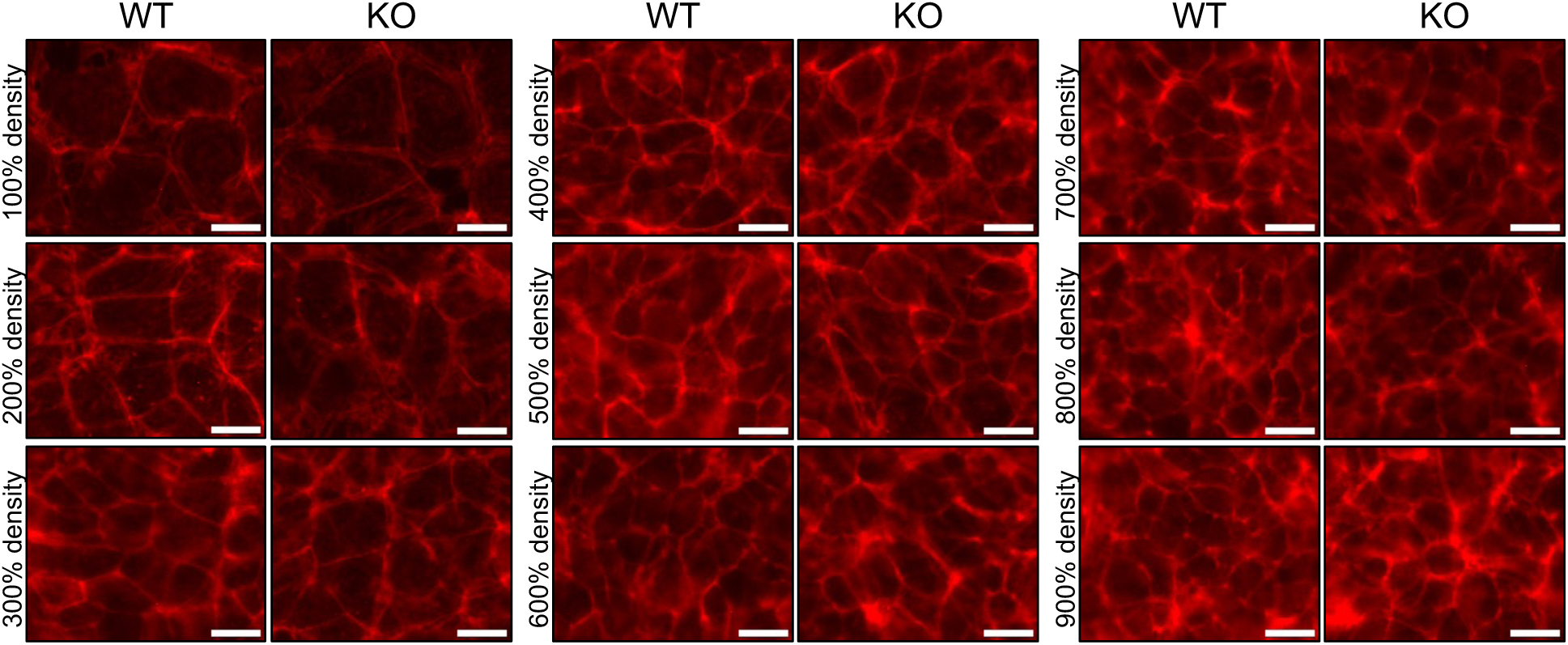
Distribution of actin filaments and cell morphology in WT and HINT1 KO cells. WT and HINT1 KO cells were cultured at various cell densities. After 24 hours, the cells were fixed and F-actin was stained with phalloidin conjugated with 568. Scale: 20μm.

**Fig. S6.**
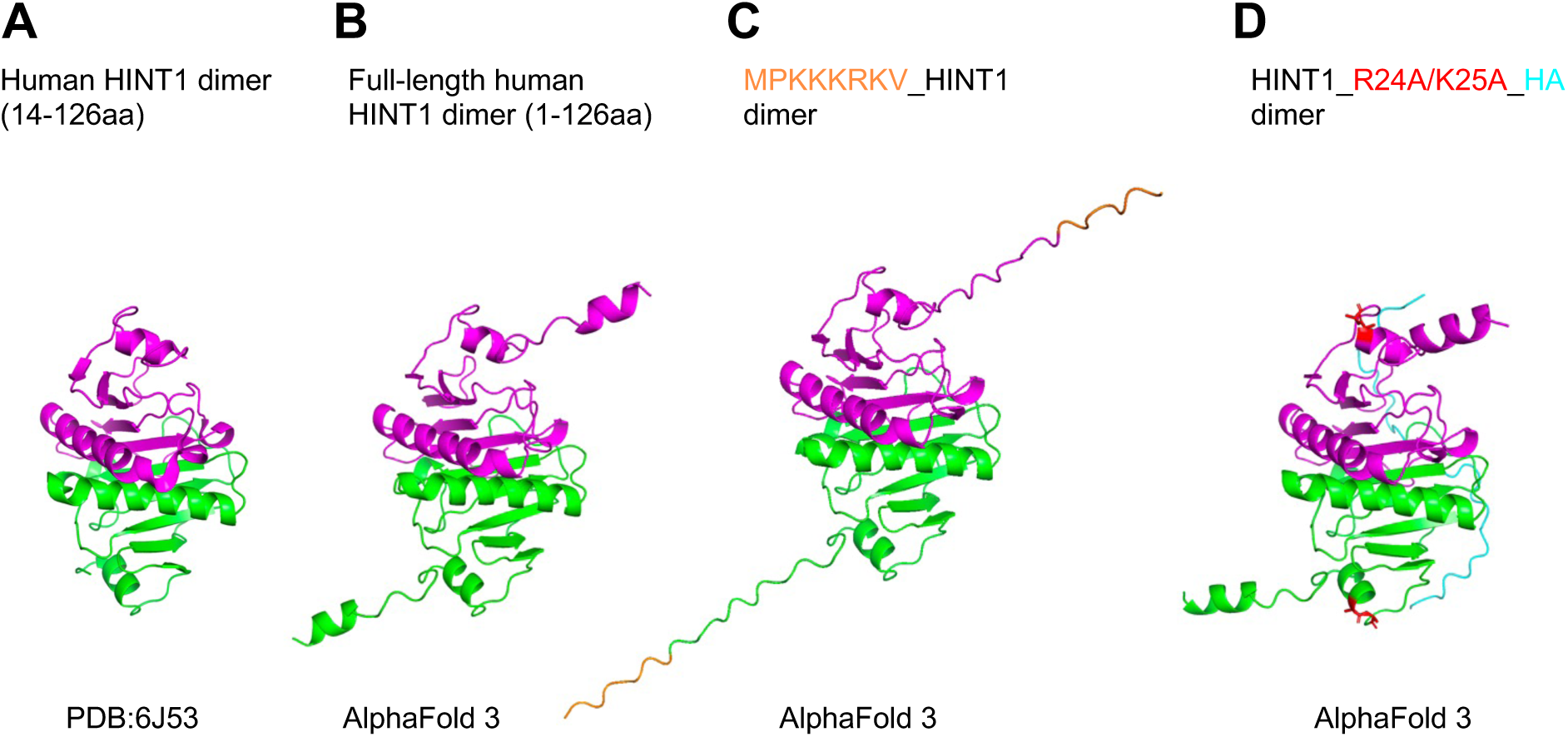
Structure of human HINT1 (A) Crystal structure of human HINT1 dimer (PDB: 6J53). Each subunit is indicated in green and magenta. Note that the N-terminal 1-13 amino acid residues are missing. (B) Structure of full-length human HINT1, predicted by AlphaFold 3, shows a striking similarity to the crystal structure. (C) Structure of human HINT1 with a nuclear localization signal peptide (MPKKKRKV indicated in orange) attached to its N-terminus. (D) Structure of human HINT1 with HA-tag. The HA-tag is indicated in cyan. KIIRKEIP(21- 28aa) is predicted as a nuclear localization signal sequence in HINT1 by nuclear localization signal prediction tool (http://www.moseslab.csb.utoronto.ca/NLStradamus/, Prediction Cutoff=0.05). R24 and K25 are mutated to Ala (A) and are indicated in red.

**Fig. S7.**
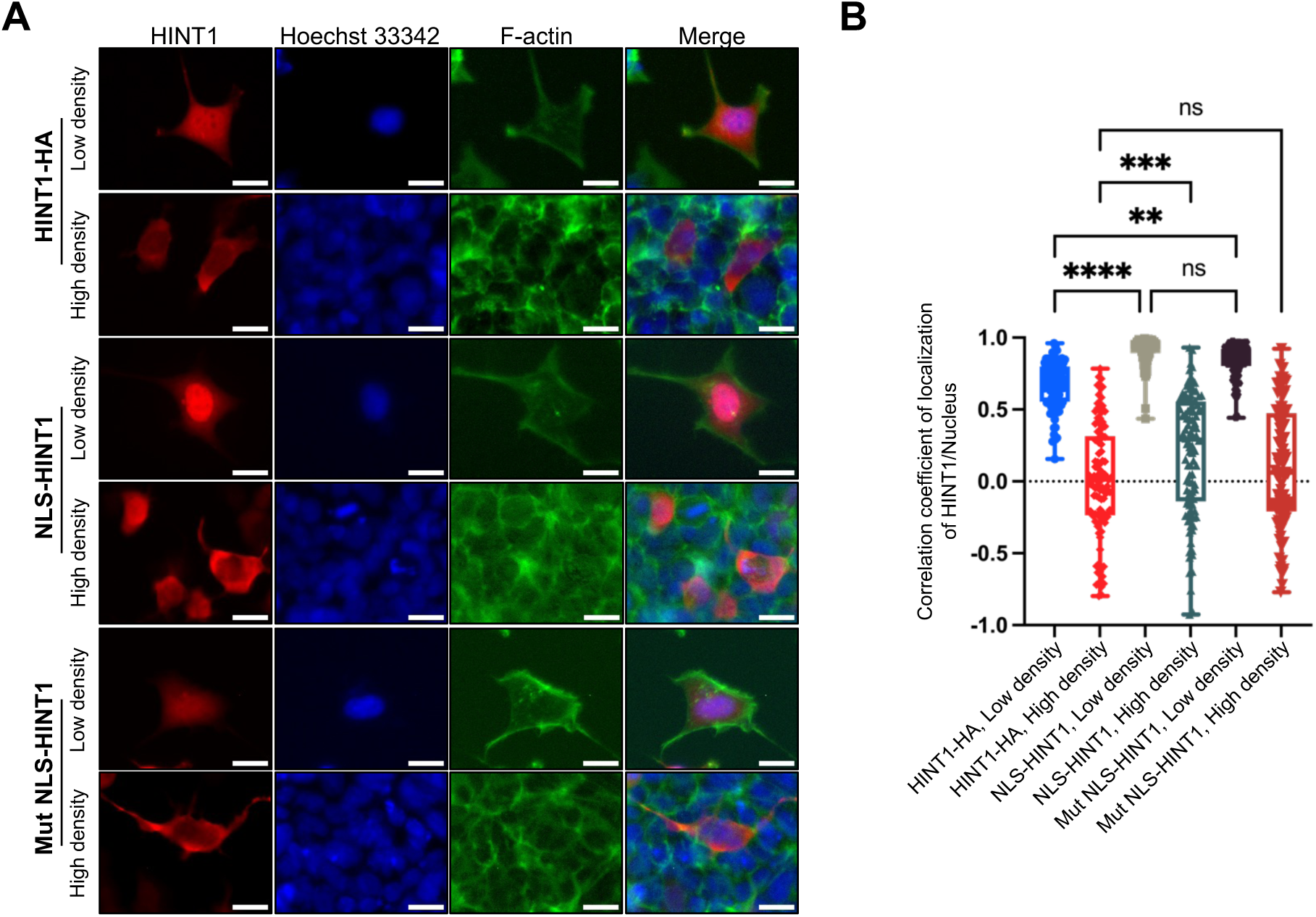
Attachment of NLS (MPKKKRKV) promote localization of HINT1 in the nucleus of low density cells but did not change its cytoplasmic localization at high density (A) NLS (MPKKKRKV-) or mutant NLS (MPKTKRKV) was fused to N-terminal of HINT1 and exogenously expressed in HEK293A HINT1-KO cells. The cells were stained as shown in Fig. 1D. Scale: 20 μm. (B) Correlation coefficient of HINT1 staining and Hoechst 33342 staining were calculated and plotted. Note that attaching an NLS promotes HINT1 localization to the nucleus, but it still remains in the cytoplasm at high cell density. Additionally, mutating the NLS does not affect the localization of HINT1.

**Fig. S8.**
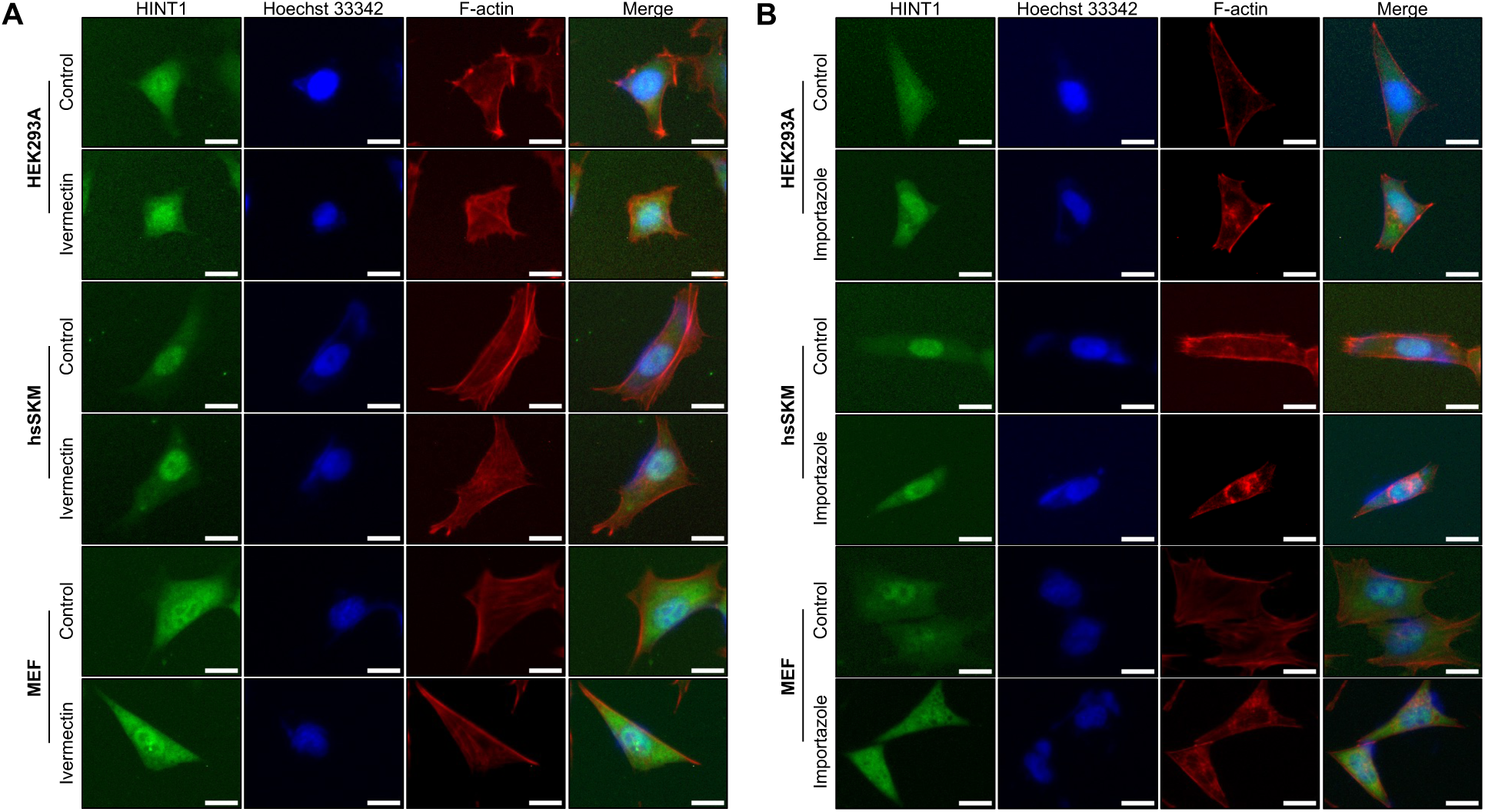
The translocation of HINT1 to the nucleus is not inhibited by importin inhibitors (A) Cells (HEK293A, hsSKM, and MEF) were treated with or without 10 μM ivermectin, an importin α/β1 complex inhibitor, for 6 hours, followed by staining as shown in Fig. 6A. (B) Cells (HEK293A, hsSKM, and MEF) were treated with or without 20 μM importazole, an importin β inhibitor, for 24 hours at low density, and then stained as shown in Fig. 6A. Scale: 20 μm.

**Fig. S9.**
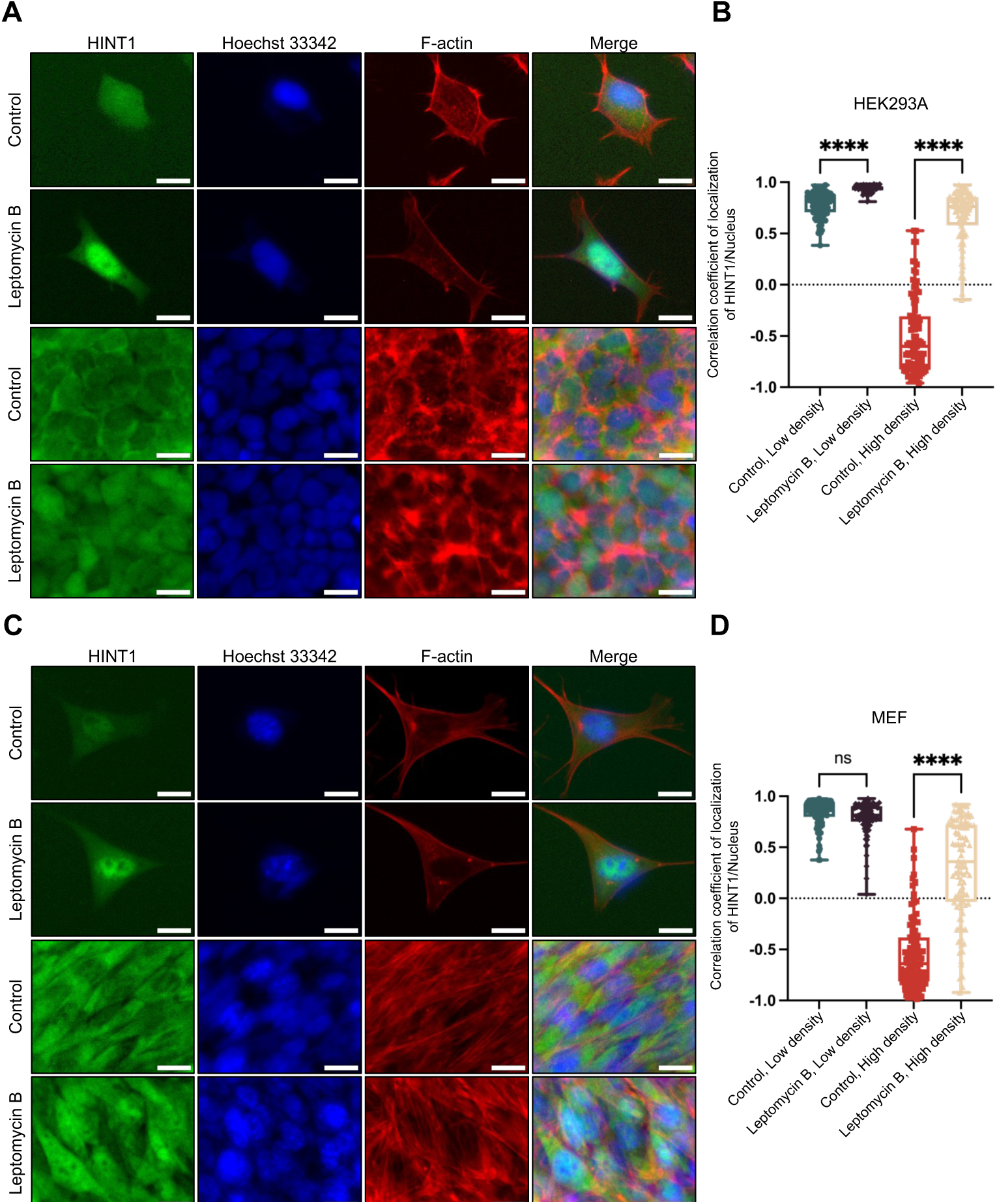
The translocation of HINT1 to the cytoplasm is inhibited by Leptomycin B, an inhibitor of exportin1/CRM1, at both low and high cell densities Cells (**A**, HEK293A; **C**, MEF cells) were treated with or without 10 nM of Leptomycin B for 24 hours at both low and high densities, then stained as shown in Fig. 6A. Correlation coefficient of HINT1 staining and Hoechst nuclear staining were calculated and plotted (**B**, HEK293A; **D**, MEF cells). Scale: 20 μm.

## References and Notes

1. Stramer, B. & Mayor, R. (2017) Mechanisms and in vivo functions of contact inhibition of locomotion, Nat Rev Mol Cell Biol. 18, 43–55.

2. Nakamura, F. (2024) The Role of Mechanotransduction in Contact Inhibition of Locomotion and Proliferation, Int J Mol Sci. 25.

3. Puliafito, A., Hufnagel, L., Neveu, P., Streichan, S., Sigal, A., Fygenson, D. K. & Shraiman, B. I. (2012) Collective and single cell behavior in epithelial contact inhibition, Proc Natl Acad Sci U S A. 109, 739–44.

4. Rubin, R. W., Warren, R. H., Lukeman, D. S. & Clements, E. (1978) Actin content and organization in normal and transformed cells in culture, J Cell Biol. 78, 28–35.

5. Hormia, M., Badley, R. A., Lehto, V. P. & Virtanen, I. (1985) Actomyosin organization in stationary and migrating sheets of cultured human endothelial cells, Exp Cell Res. 157, 116–26.

6. Toh, B. H., Yildiz, A., Sotelo, J., Osung, O., Holborow, E. J. & Fairfax, A. (1979) Distribution of actin and myosin in muscle and non-muscle cells, Cell Tissue Res. 199, 117–26.

7. Bereiter-Hahn, J. & Kajstura, J. (1988) Scanning microfluorometric measurement of TRITC-phalloidin labelled F-actin. Dependence of F-actin content on density of normal and transformed cells, Histochemistry. 90, 271–6.

8. Kajstura, J. & Bereiter-Hahn, J. (1989) Loss of focal contacts accompanies the density dependent inhibition of cell growth, Cell Biol Int Rep. 13, 377–83.

9. Jou, T. S. & Nelson, W. J. (1998) Effects of regulated expression of mutant RhoA and Rac1 small GTPases on the development of epithelial (MDCK) cell polarity, J Cell Biol. 142, 85–100.

10. Li, Y., Kong, F., Jin, C., Hu, E., Shao, Q., Liu, J., He, D. & Xiao, X. (2019) The expression of S100A8/S100A9 is inducible and regulated by the Hippo/YAP pathway in squamous cell carcinomas, BMC Cancer. 19, 597.

11. Knight, J. B., Yamauchi, K. & Pessin, J. E. (1995) Divergent insulin and platelet-derived growth factor regulation of focal adhesion kinase (pp125FAK) tyrosine phosphorylation, and rearrangement of actin stress fibers, J Biol Chem. 270, 10199–203.

12. Wassler, M. J. & Shur, B. D. (2000) Clustering of cell surface (beta)1,4-galactosyltransferase I induces transient tyrosine phosphorylation of focal adhesion kinase and loss of stress fibers, J Cell Sci. 113 **Pt** **2**, 237–45.

13. Koyama, Y., Fukuda, T. & Baba, A. (1996) Inhibition of vanadate-induced astrocytic stress fiber formation by C3 ADP-ribosyltransferase, Biochem Biophys Res Commun. 218, 331–6.

14. Nobes, C. D., Lauritzen, I., Mattei, M. G., Paris, S., Hall, A. & Chardin, P. (1998) A new member of the Rho family, Rnd1, promotes disassembly of actin filament structures and loss of cell adhesion, J Cell Biol. 141, 187–97.

15. McDonald, J. R. & Walsh, M. P. (1985) Ca2+-binding proteins from bovine brain including a potent inhibitor of protein kinase C, Biochem J. 232, 559–67.

16. Robinson, K. & Aitken, A. (1994) Identification of a new protein family which includes bovine protein kinase C inhibitor-1, Biochem J. 304 ( **Pt 2**), 662–4.

17. Dillenburg, M., Smith, J. & Wagner, C. R. (2023) The Many Faces of Histidine Triad Nucleotide Binding Protein 1 (HINT1), ACS Pharmacol Transl Sci. 6, 1310–1322.

18. Genovese, G., Ghosh, P., Li, H., Rettino, A., Sioletic, S., Cittadini, A. & Sgambato, A. (2012) The tumor suppressor HINT1 regulates MITF and beta-catenin transcriptional activity in melanoma cells, Cell Cycle. 11, 2206–15.

19. Li, K., Li, Y. & Nakamura, F. (2023) Identification and partial characterization of new cell density-dependent nucleocytoplasmic shuttling proteins and open chromatin, Sci Rep. 13, 21723.

20. Aragona, M., Panciera, T., Manfrin, A., Giulitti, S., Michielin, F., Elvassore, N., Dupont, S. & Piccolo, S. (2013) A mechanical checkpoint controls multicellular growth through YAP/TAZ regulation by actin-processing factors, Cell. 154, 1047–1059.

21. Dupont, S., Morsut, L., Aragona, M., Enzo, E., Giulitti, S., Cordenonsi, M., Zanconato, F., Le Digabel, J., Forcato, M., Bicciato, S., Elvassore, N. & Piccolo, S. (2011) Role of YAP/TAZ in mechanotransduction, Nature. 474, 179–83.

22. Li, Y., Li, K. & Nakamura, F. (2024) Characterization of open chromatin sensitive to actin polymerization and identification of core-binding factor subunit beta as mechanosensitive nucleocytoplasmic shuttling protein, Cytoskeleton (Hoboken).

23. Morel, V., Campana-Salort, E., Boyer, A., Esselin, F., Walther-Louvier, U., Querin, G., Latour, P., Lia, A. S., Magdelaine, C., Beze-Beyrie, P., Behin, A., Delague, V., Levy, N., Stojkovic, T., Attarian, S. & Bonello-Palot, N. (2022) HINT1 neuropathy: Expanding the genotype and phenotype spectrum, Clin Genet. 102, 379–390.

24. Suchanecka, A., Boron, A., Chmielowiec, K., Stronska-Pluta, A., Masiak, J., Lachowicz, M., Chmielowiec, J., Trybek, G. & Grzywacz, A. (2024) HINT1 Gene Polymorphisms, Smoking Behaviour, and Personality Traits: A Haplotype Case-Control Study, Int J Mol Sci. 25.

25. Kalderon, D., Roberts, B. L., Richardson, W. D. & Smith, A. E. (1984) A short amino acid sequence able to specify nuclear location, Cell. 39, 499–509.

26. Jans, D. A., Martin, A. J. & Wagstaff, K. M. (2019) Inhibitors of nuclear transport, Curr Opin Cell Biol. 58, 50–60.

27. Garzon-Nino, J., Rodriguez-Munoz, M., Cortes-Montero, E. & Sanchez-Blazquez, P. (2017) Increased PKC activity and altered GSK3beta/NMDAR function drive behavior cycling in HINT1-deficient mice: bipolarity or opposing forces, Sci Rep. 7, 43468.

28. Hartwig, J. H., Thelen, M., Rosen, A., Janmey, P. A., Nairn, A. C. & Aderem, A. (1992) MARCKS is an actin filament crosslinking protein regulated by protein kinase C and calcium-calmodulin, Nature. 356, 618–622.

29. Calabrese, B. & Halpain, S. (2005) Essential role for the PKC target MARCKS in maintaining dendritic spine morphology, Neuron. 48, 77–90.

30. Graff, J. M., Stumpo, D. J. & Blackshear, P. J. (1989) Characterization of the phosphorylation sites in the chicken and bovine myristoylated alanine-rich C kinase substrate protein, a prominent cellular substrate for protein kinase C, J Biol Chem. 264, 11912–9.

31. Chou, T. F., Tikh, I. B., Horta, B. A., Ghosh, B., De Alencastro, R. B. & Wagner, C. R. (2007) Engineered monomeric human histidine triad nucleotide-binding protein 1 hydrolyzes fluorogenic acyl-adenylate and lysyl-tRNA synthetase-generated lysyl-adenylate, J Biol Chem. 282, 15137–47.

32. Razin, E., Zhang, Z. C., Nechushtan, H., Frenkel, S., Lee, Y. N., Arudchandran, R. & Rivera, J. (1999) Suppression of microphthalmia transcriptional activity by its association with protein kinase C-interacting protein 1 in mast cells, J Biol Chem. 274, 34272–6.

33. Weiske, J. & Huber, O. (2005) The histidine triad protein Hint1 interacts with Pontin and Reptin and inhibits TCF-beta-catenin-mediated transcription, J Cell Sci. 118, 3117–29.

34. Wang, L., Li, H., Zhang, Y., Santella, R. M. & Weinstein, I. B. (2009) HINT1 inhibits beta-catenin/TCF4, USF2 and NFkappaB activity in human hepatoma cells, Int J Cancer. 124, 1526–34.

35. Su, T., Suzui, M., Wang, L., Lin, C. S., Xing, W. Q. & Weinstein, I. B. (2003) Deletion of histidine triad nucleotide-binding protein 1/PKC-interacting protein in mice enhances cell growth and carcinogenesis, Proc Natl Acad Sci U S A. 100, 7824–9.

36. Li, H., Zhang, Y., Su, T., Santella, R. M. & Weinstein, I. B. (2006) Hint1 is a haplo-insufficient tumor suppressor in mice, Oncogene. 25, 713–21.

37. Seburn, K. L., Morelli, K. H., Jordanova, A. & Burgess, R. W. (2014) Lack of neuropathy-related phenotypes in hint1 knockout mice, J Neuropathol Exp Neurol. 73, 693–701.

38. Sun, L., Liu, P., Liu, F., Zhou, Y., Chu, Z., Li, Y., Chu, G., Zhang, Y., Wang, J. & Dang, Y. H. (2017). Effects of Hint1 deficiency on emotional-like behaviors in mice under chronic immobilization stress, Brain Behav. 7, e00831.

39. Brzoska, P. M., Chen, H., Levin, N. A., Kuo, W. L., Collins, C., Fu, K. K., Gray, J. W. & Christman, M. F. (1996) Cloning, mapping, and in vivo localization of a human member of the PKCI-1 protein family (PRKCNH1), Genomics. 36, 151–6.

40. Klein, M. G., Yao, Y., Slosberg, E. D., Lima, C. D., Doki, Y. & Weinstein, I. B. (1998) Characterization of PKCI and comparative studies with FHIT, related members of the HIT protein family, Exp Cell Res. 244, 26–32.

41. Lima, C. D., Klein, M. G., Weinstein, I. B. & Hendrickson, W. A. (1996) Three-dimensional structure of human protein kinase C interacting protein 1, a member of the HIT family of proteins, Proc Natl Acad Sci U S A. 93, 5357–62.

42. Elosegui-Artola, A., Andreu, I., Beedle, A. E. M., Lezamiz, A., Uroz, M., Kosmalska, A. J., Oria, R., Kechagia, J. Z., Rico-Lastres, P., Le Roux, A. L., Shanahan, C. M., Trepat, X., Navajas, D., Garcia-Manyes, S. & Roca-Cusachs, P. (2017) Force Triggers YAP Nuclear Entry by Regulating Transport across Nuclear Pores, Cell. 171, 1397–1410 e14.

43. Feng, M., Wang, J., Li, K. & Nakamura, F. (2023) UBE2A/B is the trans-acting factor mediating mechanotransduction and contact inhibition, Biochem J. 480, 1659–1674.

44. Larsson, C. (2006) Protein kinase C and the regulation of the actin cytoskeleton, Cell Signal. 18, 276–84.

45. Ringvold, H. C. & Khalil, R. A. (2017) Protein Kinase C as Regulator of Vascular Smooth Muscle Function and Potential Target in Vascular Disorders, Adv Pharmacol. 78, 203–301.

46. Enomoto, A., Murakami, H., Asai, N., Morone, N., Watanabe, T., Kawai, K., Murakumo, Y., Usukura, J., Kaibuchi, K. & Takahashi, M. (2005) Akt/PKB regulates actin organization and cell motility via Girdin/APE, Dev Cell. 9, 389–402.

47. Wu, X. S., Bao, T. H., Ke, Y., Sun, D. Y., Shi, Z. T., Tang, H. R. & Wang, L. (2016) Hint1 suppresses migration and invasion of hepatocellular carcinoma cells in vitro by modulating girdin activity, Tumour Biol. 37, 14711–14719.

48. Zhang, Y., Da, Q., Cao, S., Yan, K., Shi, Z., Miao, Q., Li, C., Hu, L., Sun, S., Wu, W., Wu, L., Chen, F., Wang, L., Gao, Y., Huang, Z., Shao, Y., Chen, H., Wei, Y., Chen, F., Han, Y., Xie, L. & Ji, Y. (2021) HINT1 (Histidine Triad Nucleotide-Binding Protein 1) Attenuates Cardiac Hypertrophy Via Suppressing HOXA5 (Homeobox A5) Expression, Circulation. 144, 638–654.

49. Feng, Z., Mao, Z., Yang, Z., Liu, X. & Nakamura, F. (2023) The force-dependent filamin A-G3BP1 interaction regulates phase-separated stress granule formation, J Cell Sci. 136.

50. Xia, Y., Li, K., Li, J., Wang, T., Gu, L. & Xun, L. (2019) T5 exonuclease-dependent assembly offers a low-cost method for efficient cloning and site-directed mutagenesis, Nucleic Acids Res. 47, e15.

51. Stringer, C., Wang, T., Michaelos, M. & Pachitariu, M. (2021) Cellpose: a generalist algorithm for cellular segmentation, Nat Methods. 18, 100–106.

